# Inducible re-epithelialization of cancer cells increases autophagy and DNA damage: implications for breast cancer dormancy

**DOI:** 10.1101/2025.02.09.636592

**Authors:** Diana Drago Garcia, Suvendu Giri, Rishita Chattaerjee, Arturo Simoni Nieves, Maha Abedrabbo, Alessandro Genna, Mary Luz Uribe Rios, Moshit Lindzen, Arunachalam Sekar, Nitin Gupta, Noa Aharoni, Tithi Bhandari, Agalyan Mayalagu, Luisa Schwarzmüller, Nooraldeen Tarade, Rong Zhu, Harsha-Raj Mohan-Raju, Feride Karatekin, Francesco Roncato, Yaniv Eyal-Lubling, Tal Keidar, Yam Nof, Nishanth Belugali Nataraj, Karin Shira Bernshtein, Bettina Wagner, Nishanth Ulhas Nair, Neel Sanghvi, Ronen Alon, Rony Seger, Eli Pikarsky, Sara Donzelli, Giovanni Blandino, Stefan Wiemann, Sima Lev, Ron Prywes, Dalit Barkan, Oscar Rueda, Carlos Caldas, Eytan Ruppin, Yosef Shiloh, Maik Dahlhoff, Yosef Yarden

## Abstract

Epithelial lineage differentiation is pivotal to mammary gland development and it can pause metastasis of breast cancer (BC) by inducing tumor dormancy. To simulate this, we expressed epithelial genes in mesenchymal BC cells. Inducible expression of the epithelial *OVOL* genes in metastatic BC cells suppressed proliferation and migration. We found that *C1ORF116*, an OVOL’s target, is susceptible to genetic and epigenetic aberrations in BC. It is regulated by steroids and functions as a putative autophagy receptor that inhibits antioxidants like thioredoxin. Accordingly, boosting epithelialization lowered glutathione, elevated reactive oxygen species and increased both DNA oxidation and double strand breaks. Epithelialization also associated with redistribution of NRF2 and an altered interplay among p38, ATM, and the other kinases regulating the DNA damage response. Hence, hormonal regulation of OVOLs and chronic stress might permit epithelial differentiation and retard exit from dormancy, while altering redox homeostasis and permitting DNA damage accumulation, which may awaken dormant tumors.

## Introduction

Epithelial lineage determination and differentiation shape the mammary epithelium during embryonic development as well as later, during gland development (*1*). Both mesenchymal-to-epithelial transition (MET) and the reverse transition, epithelial-to-mesenchymal (EMT), are associated with normal mammary development during embryogenesis, and with cancer. The EMT program depletes epithelial features, such as cell polarity, while conferring mesenchymal features.

However, rather than dualistically switching between epithelial and mesenchymal states, cells can acquire a spectrum of E/M states (*2*). Nevertheless, the relevance of EMT and especially the hybrid states to clinical metastasis is still open (*3, 4*). Likewise, although it is clear that disseminated tumor cells (DTCs) may enter prolonged quiescence, how these cells later relapse remains incompletely understood.

Relapses of dormant metastases might occur in patients as long as 15-20 years after surgical removal of the primary tumor (*5*). Once arrested in the host parenchyma, DTCs acquire a reversible phenotype that includes reduced growth and motility, along with resistance to treatment (*6, 7*). Poor vascularization might instigate angiogenic dormancy (*8*), whereas suppressive immune cells may control immunogenic dormancy (*9, 10*). Both mechanisms likely lead to balanced proliferation and apoptosis (‘tumor mass dormancy’). Alternatively, disseminated solitary or clustered cancer cells might adopt a state of reversible growth arrest (*11*). Resolving the molecular details of entry into and escape from dormancy is limited by the relatively short duration of conventional experiments and dearth of suitable models. Early studies identified increased active p38, relative to active ERK, as a marker of tumor dormancy (*12*), and later work showed that endothelium-derived angiogenesis suppressors, such as thrombospondin-1 (TSP1), as well as phosphorylation of myosin light chain (MLC) (*13*), control dormancy. Likewise, through combined signaling by the tumor necrosis factor and interferon-gamma, tumor-specific T cells prevent angiogenesis and cell proliferation, to promote dormancy (*14*). STING (Stimulator of Interferon Genes) also acts as a suppressor of reawakened metastatic cells (*15*).

Resolving the action of specific EMT-inducing transcription factors (such as ZEB, SNAIL and TWIST) has greatly contributed to the understanding of metastasis (*16*). To similarly simulate the reciprocal transition, MET, and gain insights into tumor dormancy, we induced epithelialization of highly mesenchymal BC cells by means of doxycycline-inducible alleles of two ovo-like (OVOL) proteins. The corresponding transcriptional regulators have repeatedly been implicated in epithelial lineage differentiation (*17*) and hybrid E/M states (*18–22*). For instance, OVOLs regulate epithelial differentiation during hair formation and spermatogenesis (*23*), and mathematical models identified OVOLs as stabilizers of hybrid E/M states (*24*). According to a recent study, OVOL1 suppresses EMT by accelerating degradation of the TGF-β receptor (*25*). In addition, OVOL1 is involved in somatic cell reprogramming (*26*), whereas loss of OVOL2 confers stemness characteristics (*27*).

When overexpressed in highly aggressive, mesenchymal BC cells, OVOLs both decreased the abundamcer of EMT transcription factors and increased expression of E-cadherin and C1ORF116, a poorly characterized OVOL target. Furthermore, we found that C1ORF116 functions as a putative autophagy receptor that physically controls two redox modulating proteins, thioredoxin and GCLC (glutamate-cysteine ligase catalytic), thereby elevates reactive oxygen species (ROS). Parallelly to the inactivation of antioxidants at the protein level, OVOLs can inhibit the ability of NRF2 (nuclear factor erythroid 2-related factor 2) to transcriptionally up-regulate antioxidant proteins. Consistent with this, we found that OVOL1 alters DNA repair by controlling the kinases involved in damage response, as well as by increasing DNA oxidation and double strand breaks. These findings are reminiscent of recent studies showing that (i) DTCs fail colonizing oxidized tissues (*28*), (ii) autophagy supports survival of dormant cancer cells (*29, 30*), and (iii) dormant BC metastases keep accumulating new mutations (*31*). Taken together, our results raise the possibility that epithelialization and acquisition of the dormant state are controlled by epithelial genes, such as *OVOL1*, which is hormone-dependent, regulates redox homeostasis and permits accumulation of DNA damage.

## Results

### An in vitro 3-D model of breast cancer dormancy points towards dormancy induction by OVOL proteins

Because the OVOL transcriptional repressors have been implicated in epithelial lineage differentiation and in the establishment of hybrid E/M states (*18–22*), we assumed that their overexpression in mesenchymal BC cells will promote epithelial differentiation and simulate phenotypes sharing functional features with dormant DTCs. To test this prediction, we firstly established inducible overexpression (iOE) of OVOLs in the highly mesenchymal BC cell line MDA-MB-231, carrying mutant forms of p53, BRAF, KRAS and NF1 (Fig. S1A). Next, we adopted a well-established model of cancer dormancy based on the observation that plating dormant BC cells on a 3-D extracellular matrix, called 3D-BME, decelerates proliferation due to nuclear expression of cyclin-dependent kinases, but non-dormant cells retain robust proliferation under the same conditions (*13*). In line with these observations, cells displaying growth arrest in the 3D BME system were found to be dormant in vivo (*13, 32–34*) and the experimental model has been adopted by several laboratories, including researchers who studied dormancy of breast cancer (*35*) and in osteosarcoma (*36*).

After calibrating the minimal concentration of doxycycline (DOX) that was sufficient for long-term OVOL1/2 induction, we diluted the cells, including the control EV (empty vector) cells, in media containing Basement Membrane Extract (BME). Next, the cells were overlaid on a gelled BME layer and visualized. Notably, a previous study that made use of this 3-D system attributed to collagen, actin fibers and integrin important roles in the transition from dormancy to metastatic growth (*32*). In contrast to the actin stress fibers displayed by un-induced MDA-MB-231 cells, F-actin displayed cortical distribution in cells overexpressing either OVOL1 or OVOL2 (Fig. S1B), and the cells changed their morphology from an elongated to a more rounded appearance (Fig. S1C) characteristic to arrested, non-metastatic cells (*13*). Consistent with the observed cytoskeletal reorganization, the rapidly proliferating MDA-MB-231 cells almost fully arrested their growth in the 3-D matrix once the expression of OVOL1 or OVOL2 was induced, but growth of the empty vector cells and the DOX-untreated cells was unaffected (Figs. S1D and S1E). This demonstration that OVOLs can confer dormancy traits, in vitro, to highly aggressive BC cells propmted our subsequent efforts to resolve the mechanisms underlying epithelialization and dormancy.

### OVOL1 functions as a mammary epithelium differentiation marker regulated by growth factors, hormones and malignancy

In similarity to the SNAIL family of EMT-TFs, OVOLs are zinc-finger transcriptional repressors that bind with epigenetic regulators through N-terminal SNAG domains ((*37*); Fig. 1A). Despite homologous SNAG motifs and shared domain configurations, SNAILs repress but OVOLs elevate E-cadherin (*38*). In line with distinct roles, mRNA analysis of >50 mammary cell lines (*39*) clearly separated OVOL1/2 from SNAIL1/2 and other mesenchymal markers (Fig. S1F). Further analysis confirmed absence of OVOL1 in claudin-low/ZEB1-positive cell lines (Fig. S1G), which are enriched for EMT markers (*40*). In addition, evaluation of data retrieved from the Cancer Genome Atlas Breast Invasive Carcinoma Collection (TCGA-BRCA; 1,084 patients), well separated the OVOL family from the SNAILs and other mesenchymal markers, such as ZEB1 and TWIST2 (Fig. 1B). Aside from cell lines and patients, we analyzed OVOL1/2 in a large biobank of patient-derived tumour xenograft (PDTX) models representing the diversity of human BC (*41*) (Fig. S1H). This uncovered a similar pattern in which OVOL1/2 are expressed in many models, whereas the EMT markers are, in general, lowly expressed. Of note, in both PDTX and cell line models, OVOL1 was highly correlated to epithelial lineage markers (Fig. S1I). In conclusion, although the OVOL and SNAIL families engage DNA response elements and epigenetic modifiers through homologous motifs, they respectively belong to distinct epithelial and mesenchymal groups of genes.

**Figure 1:**
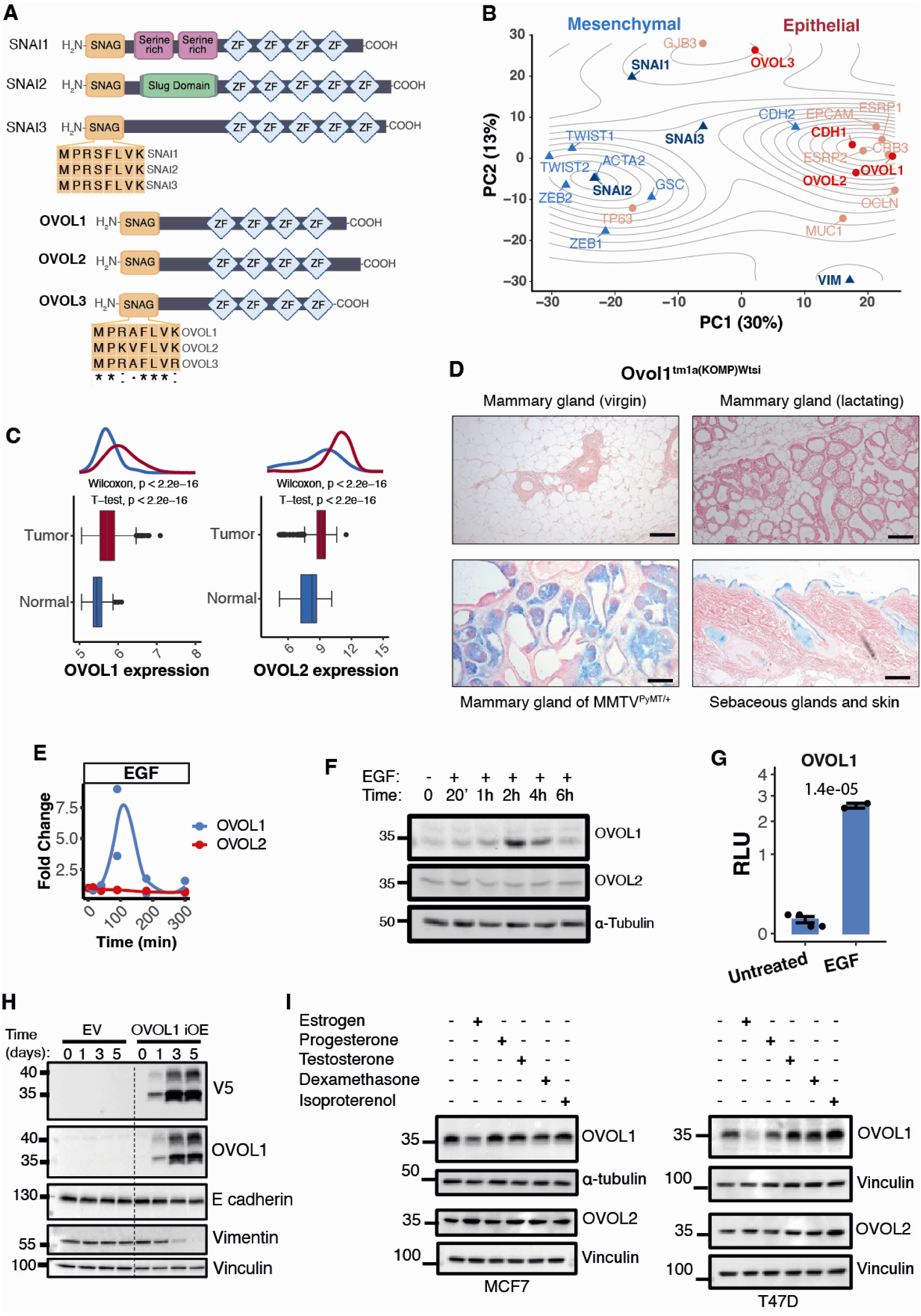
OVO-L1 is expressed in human and murine mammary cancers and acts as an EGF- and steroid hormone-inducible epithelial marker. (**A**) The domain structures of the OVO-like (OVOL) and the SNAIL (SNAI) families are shown. Note the conserved amino acid sequences of the Snail/Gfi-1 (SNAG) domains and the common configuration of the C2H2 zinc fingers (ZF) relative to the N-terminal SNAG. (**B**) Transcriptional profiles of mesenchymal (blue) and epithelial genes (red) were retrieved from the TCGA-BRCA resource (Cancer Genome Atlas Breast Invasive Carcinoma; 1,084 patients) and subjected to principal component analysis (PCA). *OVOL* and *SNAI* family members are highlighted. Contours correspond to the estimated gene density distribution. (**C**) Densities and box plots comparing the expression of OVOL1 and OVOL2 in human normal breast tissue (144 samples) and tumor samples (1980 samples; METABRIC dataset). The Welch two-sample t-test (parametric) and Mann-Whitney test (non-parametric, two-sample Wilcoxon rank-sum test) were used. Note that both tests indicate significant increase in the expression of OVOL1 and OVOL2 in tumors relative to the control samples. (**D**) Shown are beta-galactosidase-stained tissues from female mice expressing a murine *Ovol1* promoter-reporter construct. Note that the promoter is inactive in the mammary glands of virgin female mice and in lactating mammary glands (*Ovol1^tm1a(KOMP)Wtsi^*), but it displays substantial activity in sebaceous glands and in the interfollicular epidermis of the skin (*Ovol1^tm1a(KOMP)Wtsi^*), as well as in the mammary gland tumors of *MMTV^PyMT/+^*/*Ovol1^tm1a(KOMP)Wtsi^*animals. The sections were counterstained with eosin. Scale bars, 100 µm. (**E** and **F**) MCF10A cells were treated with EGF (50 ng/ml) for the indicated time intervals. OVOL1’s and OVOL2’s mRNA levels were determined using RT-PCR (E). Alternatively, cell extracts were subjected to an immunoblotting assay that employed antibodies specific to the indicated proteins (F). (**G**) OVOL1’s promoter activity was measured in HEK293T cells after 3 hours of treatment using vehicle or EGF (50 ng/ml). The luciferase luminescence signal was normalized to secreted alkaline phosphatase (SEAP) to control for transfection efficiency. Shown are the results of 4 independent transfection experiments. (**H**) Inducible overexpression (iOE) of an OVOL1 protein tagged with a V5 peptide was pre-established in MCF10A cells. These cells, along with the control empty vector (EV) cells, were treated for the indicated time intervals with the inducer, doxycycline (1µg/ml). Lysates were subjected to immunoblot analysis, as indicated. Vinculin was used to control for equal gel loading. (**I**) MCF7 and T47D cells were treated for 24 hours with estrogen, progesterone, testosterone (DHT), dexamethasone and isoproterenol (each at 100 nM). Cell extracts were analyzed using the indicated antibodies.

Analysis of METABRIC (*42*), a BC dataset (1980 patients) that includes 144 normal mammary samples, revealed that the expression levels of OVOL1 and OVOL2 are higher in tumors relative to normal mammary tissues (Fig. 1C; p<2.2e-16). This was confirmed by analyzing OVOL1 in another dataset, TCGA-BRCA (The Cancer Genome Atlas - Breast Invasive Carcinoma), which showed that OVOL1’s levels are significantly lower in normal breast tissue compared to all subtypes of breast cancer, including the basal and the luminal B class. To independently establish these observations, we examined expression of the murine form of OVOL1. An *Ovol1* promoter fused to the beta-galactosidase gene was firstly examined in *Ovol1^tm1a(KOMP)Wtsi^* transgenic mice. As previously reported (*43*), the promotor was active in sebaceous glands and in the interfollicular epidermis of the skin (Fig. 1D). However, we detected weaker or no signals in the mammary glands of both virgin and lactating mice. Next, we bred the transgenic mice with *MMTV^PyMT/+^* animals, an accepted model of invasive BC, which expresses the polyomavirus middle T oncogene in the mammary gland (*44*). All mammary gland tumors observed in the *MMTV^PyMT/+^*/*Ovol1^tm1a(KOMP)Wtsi^* animals stained positively for beta-galactosidase (Fig. 1D), suggesting that undefined oncogenic cues elevate OVOL1 expression in both human and murine mammary glands.

Common oncogenic cues might include growth factors and steroid hormones. Hence, we firstly asked if EGF can regulate OVOL1/2 expression. Treating untransformed MCF10A mammary cells with EGF revealed that OVOL1’s mRNA and protein levels peaked 1-2 hours after stimulation, but OVOL2’s levels remained undetectable (Figs. 1E and 1F), likely because OVOL11 represses OVOL2’s promoter (*45*). By means of an OVOL1’s promoter-reporter construct we confirmed the ability of EGF to up-regulate *OVOL1* transcription (Fig. 1G). Next, we utilized OVOL1’s cDNA tagged with a V5-peptide to establish MCF10A derivatives that inducibly overexpress an allele of OVOL1 (Figs. 1H). Note that tagging OVOL1 with V5 does not affect binding to DNA elements and activation of transcription (*46*). In line with epithelial characteristics, overexpression of OVOL1 was paralleled by decreased abundance of vimentin. Because MCF10A cells are devoid of the estrogen and progesterone receptors (ER and PR, respectively), we used luminal cell lines and found that estrogen and glucocorticoids can respectively down- and up-regulate the abundance of OVOL1, but OVOL2 was less responsive and other agents, including isoproterenol, which stimulates the beta-adrenergic receptor, displayed less consistent effects (Fig. 1I). In summary, OVOL1 and OVOL2 are lowly expressed in mesenchymal BC lines and in PDTXs, but expression of OVOL1, more than OVOL2, can be controlled by growth factors and hormones. Moreover, malignanct transformation elevates OVOL1 expression in the murine mammary gland.

### High OVOL1 predicts poor prognosis of patients with ER-low tumors and associates with reduced cell proliferation, migration and matrix degradation

To uncover potential clinical significance of OVOL1/2, we divided the METABRIC dataset (*42*) to two groups according to OVOL1 transcript levels. This showed only small differences in terms of patient survival time (Fig. 2A). However, when patients were grouped into ER-low and ER-high cohorts, we observed better curve separation, especially in the ER-low cohort (Fig. S2A), in line with an association between low ER, high OVOL1, and an aggressive disease. This conclusion was strengthened by similar analyses we performed with data from TCGA-BRCA, as well as by immunohistochemical (IHC) analyses of OVOL1 levels (Fig. 2B). However, similar analyses of OVOL2, which are not shown, were less informative.

**Figure 2:**
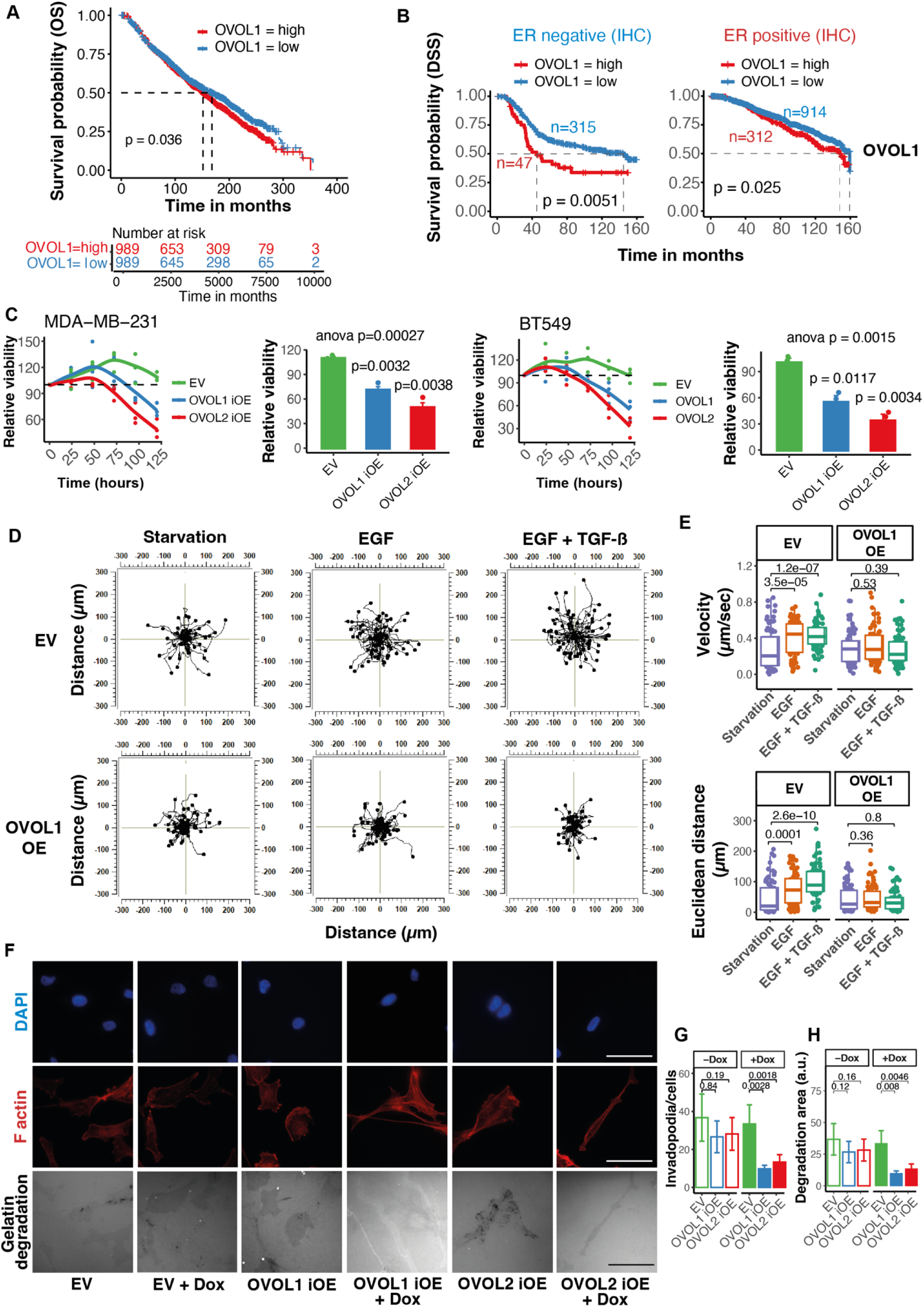
OVOL1 is associated with poor survival of patients with ER-negative BC; its overexpression reduces viability, invasiveness and migration of BC cells. (**A**) The METABRIC mRNA dataset (1980 patients) was divided to two equal groups, high (>50th percentile) and low (<50^th^) percentile, according to *OVOL1* expression level. Shown are analyses of overall patient survival per group. (**B**) The METABRIC dataset was stratified according to the status of the estrogen receptor (ER), high or low, and disease-specific patient survival (DSS) probability was calculated. The abundance of the OVOL1 protein was based on immunohistochemical (IHC) analyses. The *p* and n values are indicated. (**C**) MDA-MB-231 and BT549 cells that inducibly overexpress OVOL1 or OVOL2 were plated in 96-well plates and incubated overnight. At the indicated time points, the medium was replaced with full medium containing doxycycline (1 µg/ml). Thereafter, cells were incubated for 2 hours at 37°C with MTT (0.5 mg/ml) containing media and the formazan crystals formed by metabolically active cells were dissolved. Absorbance was determined at 570 nm. The histograms present the signals determined at the end of the 5-day interval. The experiment was performed using 3 technical replicates and 4 biological replicates. (**D** and **E**) MDA-MB-231 cells stably expressing OVOL1 or an empty vector were starved for serum factors for 20 hours. Thereafter, 40,000 cells were seeded on 0.4 µ slides (from Ibidi) that were pre-coated with fibronectin. After incubating for 2 hours to let the cells adhere, the medium was replaced with a medium containing albumin and EGF, without or with TGF-beta. Cells were imaged every 15 minutes for 6 hours. The movement and velocity of 80 individual cells were quantified and analyzed using Fiji and the Chemotaxis and Migration Tool from Ibidi. (**F** and **H**) MDA-MB-231 cells (40,000 per well; 12-well trays) were plated on Oregon-488 conjugated gelatin. The cells were untreated or treated with doxycycline (DOX; 0.25 mg/ml). Following a 24-hours long incubation, cells were fixed and stained with Alexa 568-phalloidin and DAPI. The assay used triplicates. Statistical significance of the numbers of invadopodia per cell (G) and the area of degraded gelatin (H) was analyzed using one-way Anova. *, p < 0.05; **, p<0.01; ***, p<0.005. EV, empty vector.

To permit additional analyses of OVOL1/2, we established yet another iOE derivative, of BT549 cells, which harbor partial RB gene deletions and p53 mutations. Consistent with the possibility that OVOLs confer quiescence, a distinctive hallmark of tumor dormancy (*7, 47*), DOX-induced overexpression of either OVOL gene inhibited proliferation of both BT459 and MDA-MB231 cells (Fig. S2B; note the stronger effect of cytosine arabinoside, Ara-C, a reference chemotherapeutic agent). Two additional assays were consistent with OVOL-induced growth arrest: MTT (3-(4,5-dimethylthiazol-2-yl)-2,5-diphenyltetrazolium bromide) assays, which measure mitochondrial activity (Fig. 2C), and colony formation tests (Fig. S2C).

To examine the prediction that OVOL1 can reduce cell migration, in line with the immotile phenotype of DTCs that reached dormancy (*48*), we began by establishing MDA-MB-231 cells that stably expressed OVOL1. The cells were starved for serum factors and thereafter they were placed in fibronectin-coated chambers suitable for migration assays. Notably, overexpression of OVOL1 did not change the rate of spontaneous (basal) cell migration. In contrast, the ability of EGF, either alone or in combination with the transforming growth factor beta (TGF-b), to accelerate cell migration, which was clearly manifested by the control cells, was nearly lost when OVOL1-overexpressing cells were examined (Figs. 2D and 2E). These observations implied that the upstream signals regulating cell migration, rather than the migration machinery, are controlled by OVOL1. To address matrix degradation, we assayed formation of invadopodia, protrusive structures that remodel the extracellular matrix (ECM) (*49*). DOX-inducible MDA-MB-231 cells were plated on fluorescent gelatin and after 24 hours they were fixed and stained to detect actin-rich puncta co-localizing with gelatin degradation spots. The results presented in Figures 2F-H indicated that inducible overexpression of either OVOL1 or OVOL2 significantly reduced formation of invadopodia, as well as diminished areas of matrix degradation, implying that not only inducible motility but also matrix degradation is inhibited by OVOLs.

Next, we examined in animals the effects of an overexpressed OVOL1 on tumorigenic growth and metastatic spread of MDA-MB-231 cells. Prior to implanting iOE cells in the fat pad of female mice, we either untreated or treated cells with DOX. In addition, by supplementing the animals’ drinking water with DOX, treatment of a second group of mice was continued post-implantation. A third group received DOX only after implantation. Regardless of the mode of DOX treatment, overexpression of OVOL1 strongly inhibited tumor growth (Figs. S2D and S2E). As expected, analysis of tumor extracts confirmed induction of OVOL1, along with DOX-induced increased abundance of ZEB1 and vimentin (Fig. S2F). Alongside, we observed increased abundance of two forms of C1ORF116, a transcriptional target of OVOLs (see below). Although control mice displayed more metastatic lung nodules, normalization to primary tumor weight weakened the possibility that OVOL1 markedly altered metastasis in this experiment (Figs. S2G and S2H). In conclusion, OVOL1 associates with poor prognosis of ER-low patients but its overexpression can reduce growth rates in cell cultures, as well as in a xenograft model.

### RNA sequencing and protein mass-spectrometry implicate OVOL1 in hormone response and epithelialization, as well as uncover induction of a new target gene, *C1ORF116*

RNA sequencing and proteomic analyses of MDA-MB-231 cells overexpressing OVOL1 (Fig. 3A and Supplementary Excel Files 1 and 2) concordantly uncovered increased expression of genes involved in a broad spectrum of cellular functions, such as RHOD, an atypical Rho GTPase that controls actin filaments (*50*), and SH2D3A, which is induced by hypoxia (*51*). Further analysis of the RNA data indicated that genes involved in the response to estrogen were enriched in OVOL1-overexpressing cells (Fig. S3A), whereas mesenchymal genes were enriched in the control cells (Figs. 3B and 3C). In accordance, RT-PCR confirmed decreased abundance of vimentin and increased E-cadherin in the OVOL1-overexpressing cells (Fig. 3B). Likewise, one of the characteristic epithelial genes that underwent increased abundance encodes an EGFR ligand, amphiregulin (AREG), which was verified using culture media (Fig. 3D). Additional experiments validated increased abundance of TROP2, an epithelial calcium signal transducer and an EMT-associated marker of circulating tumor cells (*52*). We also noted DOX-induced decreased and increased abundance of two additional transcripts: NR2F1 (nuclear receptor subfamily 2, group F) and C1ORF116, respectively. It has been reported that NR2F1 induces a partial EMT program and acts as a barrier to dissemination (*53*), but C1ORF116 has not previously been implicated in cancer dormancy or epithelial differentiation. We decided to focus on this poorly characterized protein because it includes no recognizable protein domain, has no family members and contains an AR (androgen receptor) response element (*54*), reminiscent of our results linking OVOLs to steroid hormones. Inducible overexpression of OVOL1/2 confirmed increased expression of two C1ORF116 forms (Fig. 3E). These forms displayed variable ratios in mammary cell lines (Fig. S3B). Along with C1ORF116, overexpression of OVOL1/2 drove increased expression of E-cadherin and KLF4, a protein implicated in epithelial differentiation (*26, 55*). Immunofluorescence analyses confirmed induction of C1ORF116 in the cytoplasm of MDA-MB-231 cells expressing an inducible allele of OVOL1 (Fig. S3C), enhanced E-cadherin and EpCAM, and decreased levels of N-cadherin, a mesenchymal marker (Figs. 3F and S3D). In summary, the uncovered transcriptional targets of OVOLs associated with increased and decreased expression of epithelial and mesenchymal markers, respectively. Notably, because OVOLs act as transcriptional repressors, the observed increased abundance of C1ORF116 likely represents a secondary effect.

**Figure 3:**
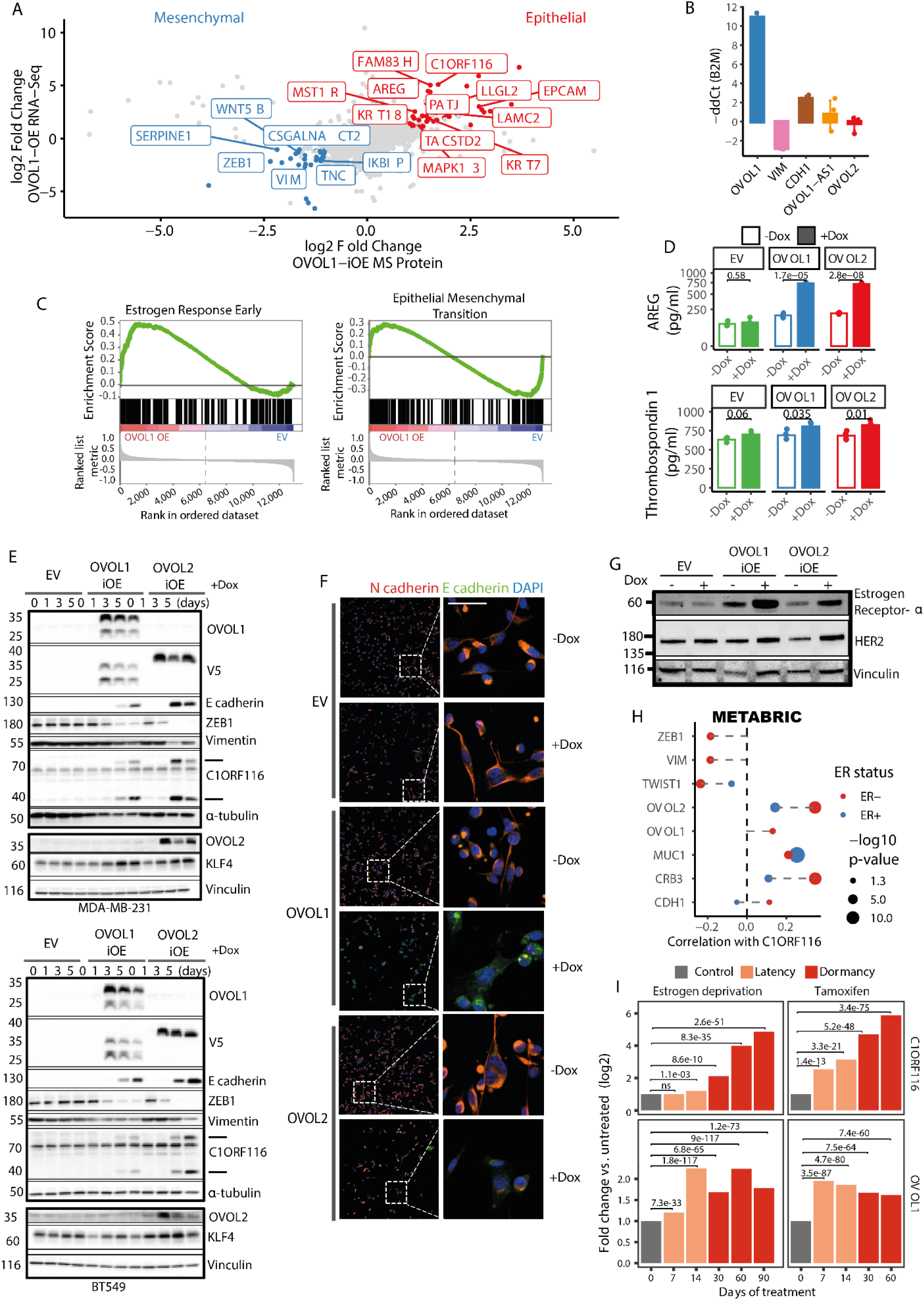
Overexpression of OVOL1 likely associates with induction of the epithelial lineage and breast cancer dormancy. (**A**) RNA and proteins were isolated from control MDA-MB-231 cells and derivatives overexpressing OVOL1. Shown are the results of both RNA sequencing analysis (log2 fold change of at least +/-1, an adjusted p-value of < 0.05, and a base mean > 5) and mass-spectrometry analysis of extracted proteins (K-nearest neighbor algorithm, KMN). The abundance of genes/proteins indicated in red are significantly increased, whereas genes/proteins colored in blue are significantly downregulated in OVOL1-overexpressing cells. The names of relevant upregulated and downregulated genes are framed in the plot. (**B**) qPCR was applied on RNA isolated from control and OVOL1-overexpressing MDA-MB-231 cells. The experiment was performed three times with 4 technical replicates. (**C**) Shown are gene set enrichment analyses (GSEA) and the enriched hallmarks that were derived from the RNA-seq data. Note that “Estrogen Response Early” was significantly enriched in the OVOL1 overexpressing cells (left panel), and “Epithelial to Mesenchymal Transition” was significantly enriched in the Empty Vector (EV) control cells (right panel). (**D**) Media conditioned for 3 days by MDA-MB-231 cells that were treated, or not, with DOX (1 µg/ml) were subjected to an ELISA specific to AREG. The results are summarized in histograms. The experiment was performed three times in triplicates. (**E**) Control MDA-MB-231 and BT549 cells (EV), along with the corresponding DOX-inducible overexpressing cells (OVOL1-iOE and OVOL2-iOE) were treated with DOX (1 µg/ml) for the indicated time intervals. Cleared cell extracts were subjected to immunoblotting that used the indicated antibodies. Anti-vinculin antibodies were used to ensure equal gel loading. (**F**) Shown are the results of immunostaining for E-cadherin and N-cadherin in OVOL1-iOE, OVOL2-iOE and control (EV) cells. Cells were treated with DOX (1 µg/ml) for 72 hours and later they were seeded on glass slides and incubated for 72 additional hours. At the end of the incubation, cells were fixed, probed using the indicated antibodies and imaged using a spinning disk microscope. Scale bar, 50 μm. Note that the framed areas shown in the left column are enlarged in the right column. (**G**) The indicated iOE derivatives of MDA-MB-231 cells, including the control empty vector (EV) cells, were treated for 72 hours with the inducer, doxycycline (DOX; 1µg/ml). Cleared cell lysates were subjected to immunoblot analysis using the indicated antibodies. Vinculin was used to control for gel loading. (**H**) RNA expression data derived from the METABRIC database of patients with BC was analyzed for correlation between C1ORF116 and expression of the indicated transcripts. Patient data were stratified according to ER status. Correlations with a *p*-values >0.05 are not shown. Correlation scores corresponding to ER-negative tumors are shown in red, and correlations corresponding to ER-positive tumors are shown in blue. (**I**) Bulk RNA sequencing data from Rosano et al., (*57*) were analyzed for the abundance of transcripts corresponding to OVOL1 and C1ORF116. Rosano and colleagues employed two MCF7 treatment models of drug-induced cancer dormancy: TAM (tamoxifen) treatment and treatment with aromatase inhibitors (estrogen deprivation). Messenger RNA was harvested at the indicated time intervals and subjected to bulk sequencing. Transcript levels were compared to the untreated arm. Note that latency refers to the time between treatment onset and dormancy entry. The data were analyzed using DESeq2. **, p<0.01; ***, p<0.005; ****, p<0.0001.

### Oncogenic RAS mutants downregulate OVOL1, expression of which is associated with C1ORF116 and both are elevated in models of cancer dormancy

To further investigate the ability of OVOLs to promote an epithelial phenotype, we examined the prediction that an oncogenic RAS mutant, widely known as a promoter of mesenchymal phenotypes (*56*), can downregulate OVOLs in human mammary cells. To this end, we stably expressed a mutant allele of HRAS in MCF10A cells and confirmed, using PCR, that the epithelial markers E-cadherin and p63 were downregulated, whereas the mesenchymal markers ZEB1, vimentin and fibronectin were upregulated (Fig. S3E). As predicted, we observed HRAS-induced increased abundance of both OVOL1 and OVOL2, in line with their inferred roles as epithelialization genes. Two additional lines of evidence lent support to the ability of OVOLs to bias epithelialization: (i) DOX-induced OVOL1 and OVOL2 increased expression of two of the major markers of the epithelial phenotype, HER2 and ER (Fig. 3G), and (ii) analysis of RNA expression data from METABRIC validated positive association between OVOLs and C1ORF116, as opposed to negative association with several epithelial markers (Fig. 3H).

To examine potential associations between cancer dormancy and both OVOL1 and C1ORF116, we analyzed a dataset published by Rosano and colleagues (*57*). These authors proposed that endocrine therapies (ETs) can induce nongenetic cell state transitions into dormancy. To demonstrate this, they expanded barcoded MCF7 cells and created multiple replicates, which were assigned to long-term estrogen deprivation (−E2; by means of a simulating treatment with aromatase inhibitors, AIs) or tamoxifen treatment (TAM). Critically, both drug-treated groups, TAM and −E2, entered a period of presumed dormancy after a few weeks of treatment, and they were maintained under pharmacological pressure until suspected awakening (early progression). We re-analyzed the data from Rosano and colleagues and summarized the results in Figure 3I. The figure depicts the average transcript levels of OVOL1 and C1ORF116 in the −E2 replicates and in the TAM replicates. Notably, both transcripts exhibited significant increased abundance, relative to baseline levels, during the latency period (the time between treatment initiation and the onset of dormancy). Furthermore, the levels of C1ORF116 continued to rise in both groups during dormancy and prior to the awakening event. Thus, along with supporting the functional link between OVOLs and OVOL1, these findings are consistent with the role we attribute to OVOL1 and C1ORF116 in tumor dormancy.

### C1ORF116 acts as an intrinsically disordered putative autophagy receptor

In similarity to the hormonal control of OVOL1/2, we observed increased and decreased abundance of C1ORF116’s protein bands following treatment of BC cells, especially T47D, with testosterone (or dexamethasone) and estrogen, respectively (Fig. 4A) (*54*). Structure prediction uncovered an intrinsically disordered 3-D configuration of C1ORF116, which includes three alpha-fold clusters embedded in highly conserved regions (Figs. 4B and S4A). While no recognizable protein domain was identified, our search for short motifs, which used iLIR (*58*), found three putative LC3-interacting regions (LIRs; Fig. 4C). Further analysis of extended LIRs (xLIRs) against a custom position-specific scoring matrix (PSSM) assigned to the SSYDFL sequence the highest score. Notably, LIR motif-containing proteins are involved in autophagy, a degradation process that selectively targets intracellular components (*59*). The autophagosomes, double membrane vesicles, are trafficked to lysosomes along with their payloads. This process involves several autophagy related (ATG) proteins, including yeast ATG8 (LC3 and GABARAPs in mammals) and a unique type of LIR-harboring adaptor proteins, termed autophagy receptors. Consistent with the possibility that C1ORF116 functions as an autophagy receptor, sequence alignments identified homologies between the N-terminal LIR motif of C1ORF116 and the LIRs of several autophagy receptors, including p62/SQSTM1 (Fig. S4B).

**Figure 4:**
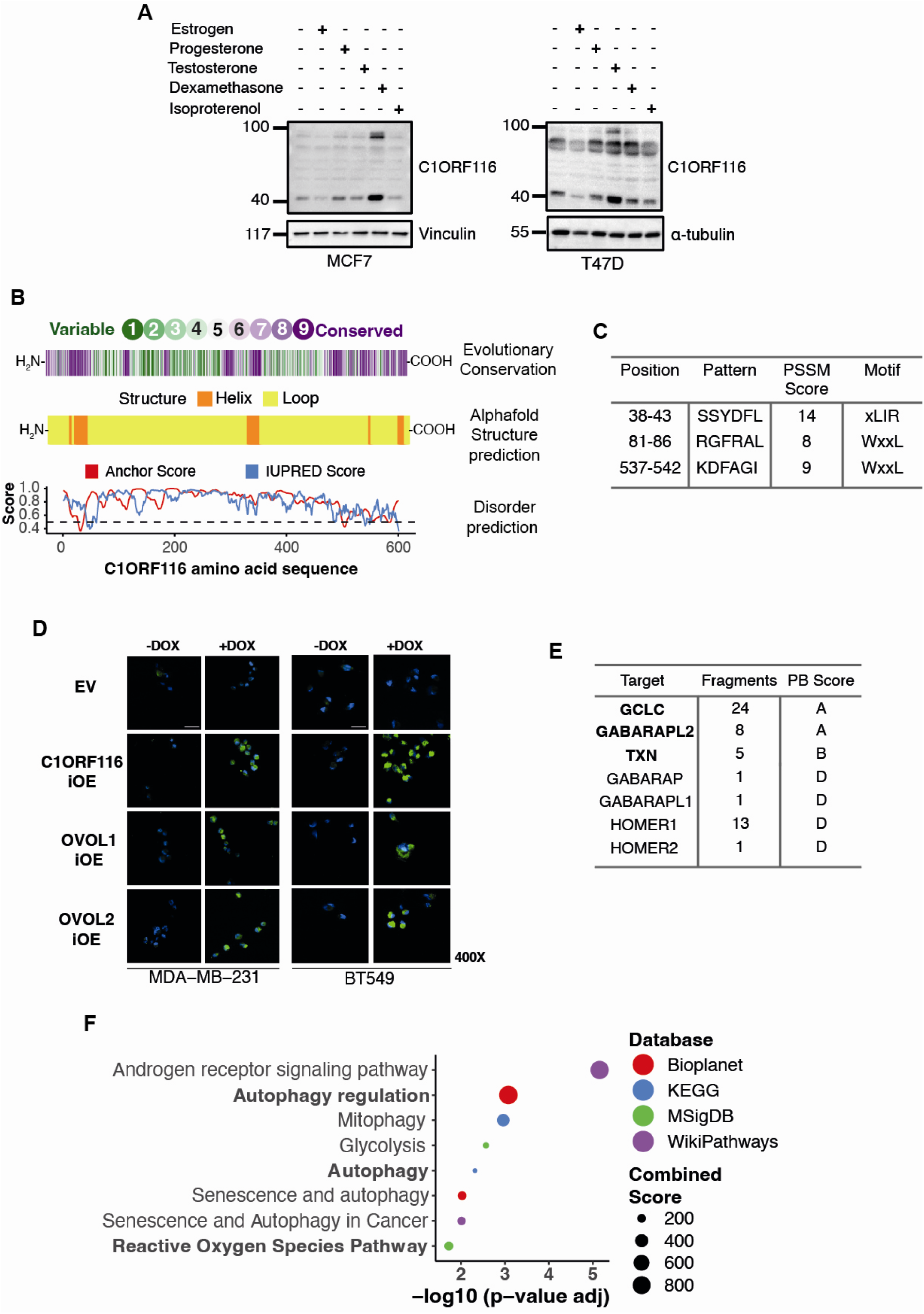
C1ORF116, a hormone-inducible intrinsically disordered protein, might act as an autophagy receptor. (**A**) MCF7 and T47D breast cancer cells were incubated for 24 hours with the indicated hormones or with isoproterenol (100 nM). Cleared cell lysates were probed for C1ORF116, as well as for a loading control protein (vinculin or tubulin). (**B**) The following parameters of C1ORF116 were analyzed: (i) Evolutionary conservation based on analysis of 73 homologous sequences (ConSurf server; https://consurf.tau.ac.il/consurf_credits.php), (ii) 3D structure, including helix (orange) and loop (yellow) regions predicted by Alphafold (https://alphafold.ebi.ac.uk/about), and (iii) prediction of disorder using two energy estimation approaches: IUPRED (blue line; https://iupred2a.elte.hu/) and Anchor (red line; http://anchor.elte.hu/). (**C**) The iLIR resource (*58*) was used for the analysis. The summary table lists the three putative LC3-interacting regions (LIRs) of C1ORF116. Also shown are the results of scoring extended LIRs (xLIRs) against a custom position-specific scoring matrix (PSSM). Note that PSSM scores of 13–17 are considered highly accurate but some verified LIRs might have scores as low as 7 (https://ilir.warwick.ac.uk/index.php). (**D**) The indicated cells were pre-incubated for 72 hours in a medium supplemented without or with DOX. Afterwards, 40,000 cells were seeded in 8-well glass bottom chambers and incubated overnight. An autophagy assay kit (from Abcam) was used to visualize cells undergoing autophagy (green). Images (400X magnification) were obtained using a spinning disk microscope. Scale bar, 50 μm. (**E**) Listed are the results of a yeast 2-hybrid screen that used a human breast tumor epithelial cell library. A fragment of C1ORF116 (amino acids 1-601) was used as the bait. The screen identified 136 positive clones among 129 million interactions. The number of prey fragments identified is indicated, along with the Predicted Biological (PB) score, in which A indicates the highest interaction confidence. (**F**) The 30 uppermost putative interaction partners of C1ORF116 from the yeast 2-hybrid screen were subjected to pathway enrichment analysis using EnrichR. The dot plot presents the corresponding GO processes. The size of each dot represents the number of genes, whereas the color of each dot represents the corresponding database we analyzed.

Since ubiquitin serves as a signal for selective autophagy and the autophagy receptor p62/SQSTM1 both binds with ubiquitin and undergoes ubiquitination (*60*), we examined the possibility that C1ORF116 can undergo ubiquitination. As predicted, probing C1ORF116 immunoprecipitates with an anti-ubiquitin antibody confirmed that both forms of C1ORF116 undergo ubiquitination (Fig. S4C). Next, we utilized ubiquitin-agarose beads that were incubated with extracts of cells that were pre-transfected with a V5-tagged C1ORF116 plasmid. In similarity to p62/SQSTM1, C1ORF116 was specifically pulled down by the immobilized ubiquitin (Figs. S4D and S4E). Because these results potentially implicated C1ORF116 in autophagy, we utilized a dye that labels autophagic vacuoles. This revealed that overexpression of either C1ORF116, OVOL1 or OVOL2 enhanced autophagy in both MDA-MB-231 and BT549 cells (Fig. 4D). In conclusion, the poorly characterized C1ORF116 protein might serve as an autophagy receptor downstream of OVOL1/2. Notably, it has been reported that autophagy prolongs BC dormancy (*61*) because it permits long-term cell survival under nutritional stress (*48, 62*).

### C1ORF116 binds with LC3/GABARAP family proteins and two key redox enzymes

To identify target proteins of C1ORF116, we performed yeast two-hybrid (Y2H) screens. The N-terminal portion of C1ORF116 was used as a bait while screening BC cDNA libraries. The screen identified 136 positive clones (Supplementary Excel File 3). The uppermost interactors identified were GABARAPL2, a member of the LC3 family, the catalytic subunit of glutamate cysteine ligase (GCLC), and TRX (thioredoxin; Fig. 4E). Notably, TRX and GCLC control the cellular redox potential by respectively acting as a major scavenger of reactive oxygen species (ROS) and an enzyme that mediates a rate-limiting step in glutathione biosynthesis (*63*). These observations are consistent with the identification of C1ORF116 as a putative autophagy receptor and indeed pathway enrichment analysis of the 30 uppermost C1ORF116’s interaction partners from the Y2H screen putatively identified autophagy, along with AR signaling, as the most relevant pathways (Fig. 4F).

To validate the results of the Y2H screens, we firstly analyzed the subcellular localization of C1ORF116, GABARAPs and GCLC. These immunofluorescence analyses indicated that all three proteins are primarily cytoplasmic (Fig. 5A). In addition, we detected areas of co-staining as well as co-localization with C1ORF116 in some puncta. Next, we performed co-immunoprecipitation experiments that used a plasmid encoding a tagged C1ORF116. The results validated specific interactions between C1ORF116 and the endogenous forms of both GABARAPL2 (Fig. 5B) and GCLC (Fig. 5C). To address TRX, we immunoprecipitated this relatively small but abundant protein from lysates of cells that were pre-transfected with increasing amounts of the V5-C1ORF116 plasmid (Fig. 5D). This confirmed the existence of TRX-C1ORF116 complexes and revealed gradual diminution of TRX. However, we detected no parallel diminution when analyzing the rather high GCLC levels. Conceivably, by acting as an autophagy receptor, C1ORF116 physically binds with both GCLC and TRX, but only the latter is detectably sorted for degradation. Assuming autophagic degradation of TRX, we compared the effects of the proteasome inhibitor, MG132, and bafilomycin A1, which prevents maturation of autophagic vacuoles. Unlike MG132-treated cells, in cells treated with bafilomycin A1 we detected a ubiquitinated form of TRX, which disappeared following overexpression of C1ORF116 (Fig. 5E). Thus, by binding with and enhancing ubiquitination of TRX, C1ORF116 likely sorts this antioxidant protein to autophagic degradation. Notably, in line with the inferred model attributing to OVOL1/2 and C1ORF116 the ability to harness autophagy and redox homeostasis, it has previously been proposed that redox mechanisms control tissue colonization by dormant BC cells (*28, 64*).

**Figure 5:**
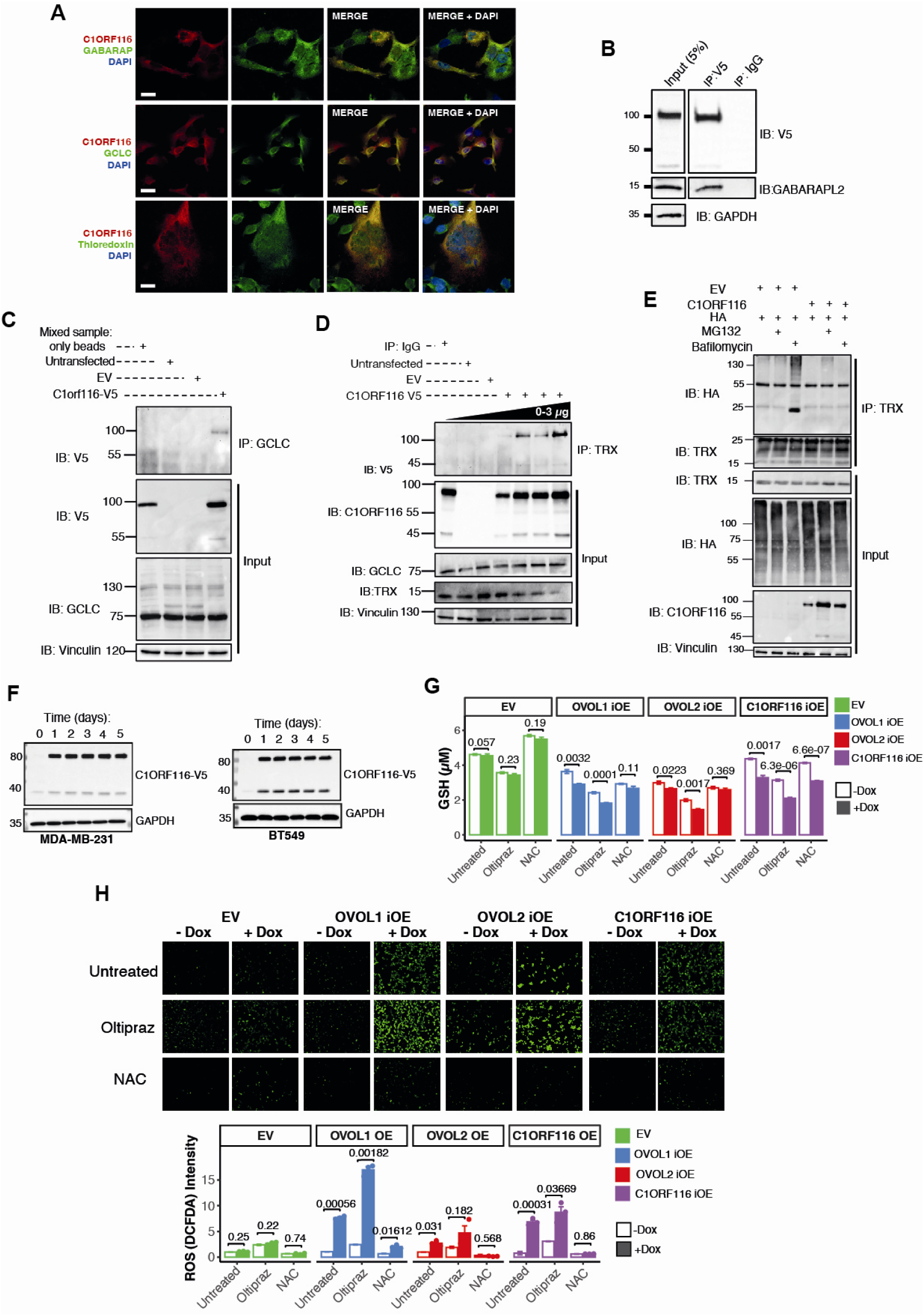
C1ORF116 binds with GABARAPL2, GCLC, and TRX, as well as increases intracellular ROS and decreases glutathione following overexpression. (**A**) MDA-MB-231 cells overexpressing an inducible V5-C1ORF116 were plated in round-bottom glass wells. After treating the cells with DOX for 3 days, cells were fixed, and immunolabelled using anti-V5 (red), anti-GABARAP (green), anti-GCLC (green) or anti-thioredoxin (green) antibodies. DAPI was used to stain nuclei (blue). Scale bar, 20 μm. (**B**) MDA-MB-231 cells stably transfected with either an empty vector (EV) or a plasmid encoding V5-C1ORF116 were lysed. The cleared extracts were subjected to immunoprecipitation (IP) using protein A/G magnetic beads coated with either normal mouse immunoglobulin G (IgG) or an anti-V5 antibody. A fraction of the lysate (5%) was separately analyzed (*Input*). After protein transfer to membranes, we performed immunoblotting (IB) with antibodies specific to the indicated proteins. (**C**) HEK293T cells grown in 90-mm dishes were un-transfected, transfected with an empty vector (EV) or with a C1ORF116-V5 plasmid. Forty-eight hours later, the cells were extracted and the cleared extracts were subjected to immunoprecipitation using protein A/G beads and antibodies specific to GCLC. As control, we mixed extracts from the EV- and the C1ORF116-transfected cells and used control beads. (**D**) HEK293T cells were transfected with an empty vector or with increasing amounts of the C1ORF116-V5 plasmid (0µg, 0.5µg, 0.75µg, 1.5µg and 3µg per plate). Forty-eight hours later, the cells were extracted and a portion of each cleared extract was subjected to immunoprecipitation (IP) with either control innunoglobulins or with a an anti-TRX antibody. Whole extracts (input) along with the immunoprecipitates were immunoblotted (IB) using the indicated antibodies, including antibodies specific to TRX and GCLC. (**E**) HEK293T cells grown in 10-cm plates were transfected with a control plasmid (EV) or with plasmids encoding HA-ubiquitin and C1ORF116-V5. Following 48 hours of incubation, cells were treated with either bafilomycin (20 nM; 16 hours) or MG132 (10 μM; 8 hours). Thereafter, we extracted all cells and immunoprecipitated TRX using agarose protein A/G beads pre-coated with an anti-TRX antibody. The washed immunoprecipitates or cleared extracts were subjected to immunoblotting using the indicated antibodies. (**F**) Inducible overexpression (iOE) of the C1ORF116 protein tagged by a V5 peptide was pre-established in both MDA-MB-231 and BT549 cells. These cells were treated for the indicated time intervals (in hours) with doxycycline (1µg/ml). Lysates were subjected to immunoblot analysis using antibodies specific to the indicated proteins. GAPDH was used to ensure equal gel loading. (**G**) The indicated derivatives of MDA-MB-231 cells were treated with doxycycline (1 µg/ml) for 72 hours to induce the overexpression of OVOL1, OVOL2, or C1ORF116. The corresponding cell extracts were subjected to an assay that determined the intracellular levels of reduced glutathione (GSH). Each column represents the mean ± SEM (triplicates) of at least three independent experiments. (**H**) Shown are results of a hydrogen peroxide assay that employed DCFDA (2,7 - dichlorofluorescin diacetate). Representative images of the inducible MDA-MB-231 cells overexpressing OVOL1, OVOL2 or C1ORF116 are presented. The images were obtained using epifluorescence microscopy (original magnification X100). Scale bar, 200 µm. The histogram presents the quantification of hydrogen peroxide levels by densitometric analysis. NAC (N-acetyl-l-cysteine; 10 mM) was used as a ROS scavenger and oltipraz (50 µM) was used to elevate ROS. Values represent mean +SEM (triplicates) of three independent experiments.

### The OVOL-C1ORF116 axis controls ROS and glutathione levels

In light of the yeast 2-hybrid screens, we predicted that overexpression of either C1ORF116 or OVOL1/2 will downregulate antioxidants and elevate ROS levels. To examine this scenario, we established inducible overexpression of C1ORF116 in MDA-MB-231 and BT549 cells (Fig. 5F). Next, we determined the levels of reduced glutathione (GSH) following treatment with DOX. Relative to the control cells, C1ORF116-overexpressing cells displayed lower basal levels of GSH, and a further reduction was observed post induction with DOX (Fig. 5G and S5A). As expected, similar analyses of OVOL1-iOE and OVOL2-iOE cells confirmed DOX-induced increased abundance of GSH (Fig. 5G). Note that these assays utilized two references: oltipraz, which elevates superoxide radicals, and N-acetyl cysteine (NAC), which quenches ROS.

In the next step, we applied a bioluminescence test that assayed hydrogen peroxide (H_2_O_2_). The assay was based on a cell-permeant fluorogenic dye, DCFDA (2,7 - dichlorofluorescin diacetate), that probes hydroxyl, peroxyl, and other ROS activities. The results shown in Figures 5H and S5B demonstrated the ability of both C1ORF116 and OVOL1 to upregulate ROS levels. Note, however, that OVOL2 exerted relatively small effects in MDA-MB-231 cells. To ensure its pivotal role in the induction of ROS, we knocked-down C1ORF116 using siRNA. As expected, pre-treatment with the specific siRNA reduced the abundance of ROS in all our iOE cell lines (Fig. S5C). However, siRNA-treated OVOL1-overexpressors still displayed substantial ROS signals, suggesting that OVOL1 can modify the redox potential through additional mechanisms. In line with this prediction, re-analysis of the RNA-seq data revealed changes in glutathione peroxidase 8 and several glutathione S-transferases, along with NCF2, which generates superoxide bursts, and cytochrome P450 1B1, which is involved in hyperoxic toxicity. In conclusion, overexpression of either C1ORF116 or OVOL1/2 elevated ROS and reduced GSH levels, in line with the ability of C1ORF116 to bind with both TRX and GCLC. Still, additional transcriptional targets of OVOLs likely regulate the redox potential of mesenchymal mammary cells once they acquire an epithelial phenotype.

### The OVOL-C1ORF116 axis controls NRF2, metabolites and antioxidants

Assuming that the effect of OVOLs on the cellular redox potential is accompanied by changes in additional metabolites, other than ROS and glutathione, we analyzed polar small molecules using mass spectrometry (Supplementary Excel File 4). The heatmap shown in Figure 6A presents the normalized abundance of selected metabolites. As expected, the metabolic effects induced by OVOLs were more varied and greater than those induced following overexpression of C1ORF116, the downstream target of OVOLs. Among the top differentially downregulated metabolites in both OVOL2- and OVOL1- overexpressing cells we identified taurine, an abundant non-essential amino acid that maintains glutathione stores, and both the oxidized and the reduced forms of glutathione. In addition, among other alterations, several amino acids were upregulated in OVOL-overexpressing cells, including glutamine and the cysteine-glycine dipeptide, a derivative of glutathione. Taken together, the observed effects of OVOLs on metabolism might complement the action of C1ORF116 on the cellular redox potential and reflect the proposed ability of autophagy to recycle amino acids in dormant cancer cells.

**Figure 6:**
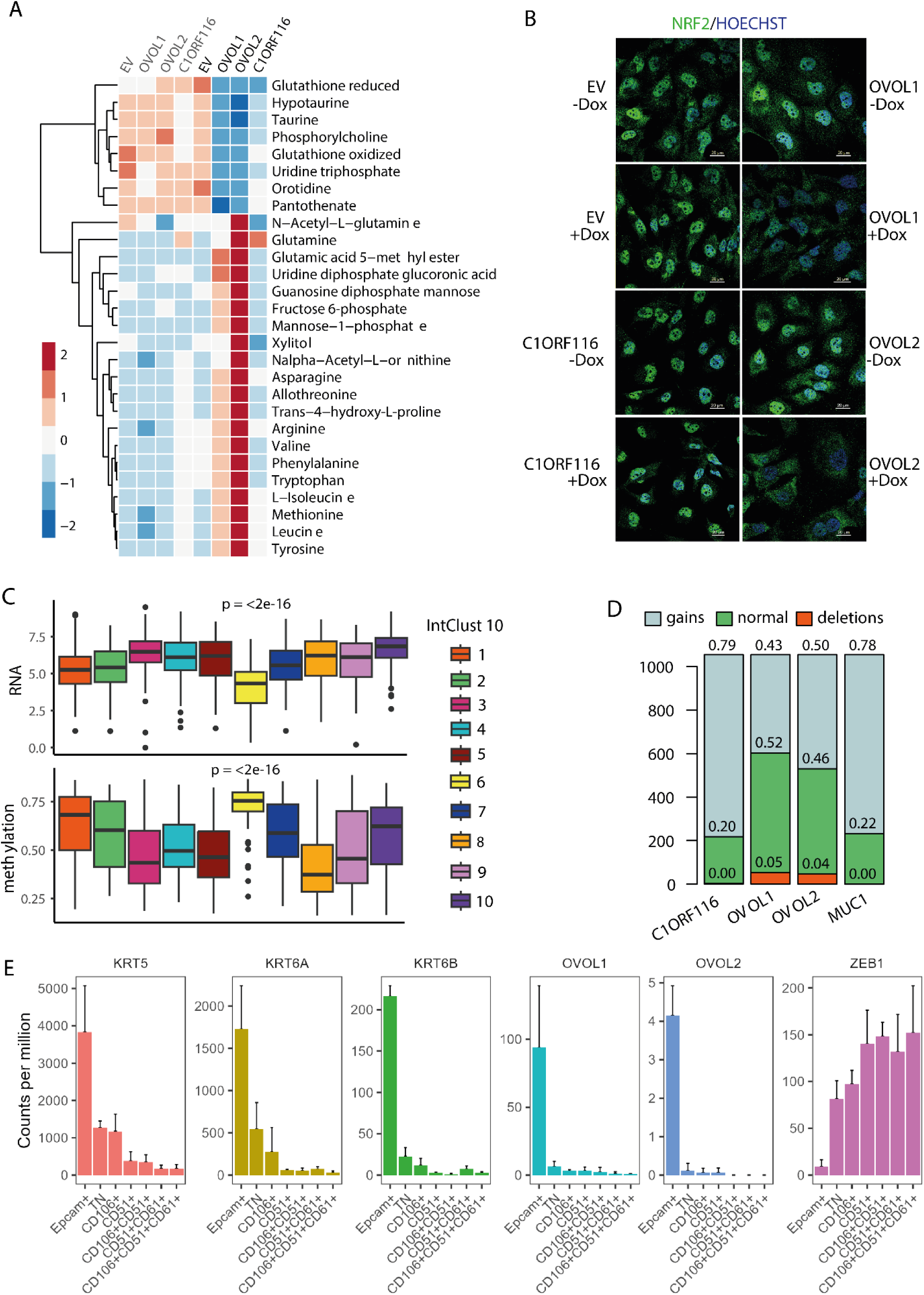
C1ORF116 displays genetic and epigenetic aberrations in BC; the upstream OVOL axis is active in epithelial E/M hybrid states and regulates both metabolism and NRF2. (**A**) Stable derivatives of MDA-MB-231 cells that overexpress OVOL1, OVOL2 or C1ORF116 (bold type), along with Empty Vector (EV; normal type) control cells, were treated for 72 hours with doxycycline or with vehicle. Thereafter, 5 million cells were snap frozen, and polar metabolites were determined using mass spectrometry. The heatmap shows the normalized abundance of selected metabolites (4 biological replicates per group). (**B**) The indicated, DOX-inducible derivatives of BT549 cells were untreated or treated with DOX for 72 hours. Thereafter, cells were counted, seeded in glass slides and treated (or not) for additional 72 hours. This was followed by fixation and confocal microscopy analysis of the subcellular distribution of NRF2 using a specific antibody. Hoechst was used to visualize nuclei. Note that NRF2 exhibits nuclear localization in control cells (EV), but it is excluded from nuclei of cells overexpressing OVOL1 or OVOL2. Bars, 20 microns. (**C**) Shown are mRNA expression levels (left panel) and methylation status (right panel) of C1ORF116 in the ten Intrinsic Cluster (ICs) of BC. ICs were derived from the invasive BC cohort of TCGA. Statistical testing of RNA and methylation levels of C1ORF116 across the IntClust 10 groups was performed using Anova. (**D**) The TCGA dataset of BC was used to derive total copy number (TCN) information for the indicated genes. Deletions were defined as TCN<2, Normal as TCN = 2, and Gains TCN>2. (**E**) Shown are levels of OVOL1 and OVOL2 expression in the previously reported stable cellular states that underlie EMT (*72*). TN refers to a triple negative state lacking expression of the epithelial cell adhesion molecule (EpCAM), as well as lacking expression of CD106 (Vascular Cell Adhesion Molecule 1; VCAM-1), CD51 (integrin alpha V; ITGAV) and CD61 (integrin beta-3; ITGB3). The data were derived from GEO record GSE110587. Error bars refer to standard error values (n=3).

The capacity of the OVOL-C1ORF116 axis to modify redox homeostasis raised the possibility that this would involve NRF2 (nuclear factor erythroid 2-related factor 2), a transcription factor capable of protecting cells against oxidative damage. Under unstressed conditions, NRF2 is unstable and retained in the cytoplasm, but in response to oxidative stress it translocates to the nucleus to activate antioxidant genes (*65*). We found that NRF2 resides mainly in the nuclei of control BT549 (Fig. 6B) and MDA-MB-231 (Fig. S5D) cells in the absence or presence of DOX. This is attributable to genetic or epigenetic aberrations characteristic of BC cells, including frequent aberrations in PTEN (*66, 67*). Nevertheless, overexpression of either OVOL1 or OVOL2 translocated NRF2 to the cytoplasm (Figs. 6B and S5D) and the following NRF2 target genes were transcriptionally repressed in OVOL1 overexpressing cells: GCLC, GSTP1, CAT and GPX4 (Fig. S5E). In contrast, NRF2 was retained in the nuclei of DOX-induced cells overexpressing C1ORF116, further implying that non-C1ORF116 mediated mechanisms contribute to the enhanced oxidative stress of OVOL-overexpressing cells. In summary, several mechanisms underlie the ability of OVOLs to enhance oxidative stress, and they include increased abundance of C1ORF116, inactivation of several antioxidants, and simultaneous suppression of the ubiquitous NRF2-centered protective machinery.

### C1ORF116 controls growth of BC cells and undergoes genetic and epigenetic aberrations in mammary tumors

It has recently been reported that C1ORF116 overexpression in thyroid cancer cells enhanced their proliferation (*68*). Consistent with this report, overexpression of C1ORF116 in MDA-MB- 231 cells accelerated proliferation, but weaker effects were observed when analyzing the less rapidly proliferating BT549 cells (Fig. S6A). Notably, whereas colony formation assays detected only small effects in both cell lines (Fig. S6B), CRISPR-CAS9 mediated ablation of *C1ORF116* in MDA-MB-231 cells clearly reduced their rate of proliferation (Fig. S6C). In line with growth stimulation, high expression levels of C1ORF116 associated, albeit weakly, with shorter survival of patients with BC (Fig. S6D). Furthermore, we noted that this prognostic effect was more significant in the group of AR-negative patients, which is reminiscent of the ability of androgens to control C1ORF116’s promoter (*54*).

To better understand the clinical significance of C1ORF116, we surveyed the 10 integrative BC clusters, which classify tumors according to their gene expression and copy number aberrations (*69*). The box plots shown in Figure 6C (upper panel) display C1ORF116 expression by each integrative cluster (IntClust). The highest level was displayed by IntClust10, which is dominated by ER negativity and multiple chromosomal aberrations. In line with the ability of estrogen to inhibit the OVOL-C1ORF116 axis, the lowest C1ORF116 level was displayed by IntClust6, comprising ER-positive/HER2-negative tumors (luminal B). Assuming that additional mechanisms can downregulate C1ORF116 in luminal B tumors, we analyzed DNA methylation profiles of >1500 breast tumors (*70*). This revealed that IntClust6 is characterized by the highest C1ORF116’s promoter methylation (Fig. 6C; lower panel), indicating that this gene is epigenetically repressed in IntClust6. In addition to aberrant methylation, we found that most tumors included in METABRIC exhibited significant copy number gains of the *C1ORF116* gene (Fig. 6D). As reference, Figure 6D also displays *MUC1*, which is one of the most aberrantly overexpressed genes in BC (*71*). According to this analysis, *C1ORF116* displays as extensive gains as *MUC1*. In conclusion, we identified C1ORF116 as a poor prognosis gene susceptible to genetic and epigenetic aberrations, as well as uncovered an association between AR negativity, high C1ORF116 and breast cancer aggressiveness.

### OVOLs are highly expressed in two stable E/T hybrid states, EpCAM-positive and TN (triple-negative)

Mathematical modeling predicts that OVOL1 stabilizes a hybrid E/M phenotype characterized by cells exhibiting both epithelial and mesenchymal traits (*24*). Accordingly, OVOL1 overexpression can reprogram fibroblasts to epithelial cells and help BC cells maintain the intermediate E/M phenotype while retarding transition to mesenchymal states (*17*). Because this mechanism could explain the poor prognosis significance of OVOLs and potential contribution to metastasis (*22*), we addressed OVOL’s relevance to the hybrid E/M states. To this end, we made use of a dataset derived from mouse tumors undergoing EMT (*72*). Originally, this analysis identified in animals several stable intermediary (hybrid) EMT states based on three surface proteins that together define all EMT transition states: CD106 (Vcam1), CD51 (ItgaV) and CD61 (Itgb3). The use of the three markers, in addotion to the epithelial cell adheion molecule, EpCAM, uncovered that OVOL1/2’s levels are highest in the fully epithelial state but they steeply decrease as the mesenchymal component increases (see Figure 6E). As references, our analysis included three basic/neutral epithelial keratins, K5, K6A and K6B, which displayed patterns similar to the distribution of OVOLs. This contrasted with ZEB1, a characteristic mesenchymal marker, which displayed gradually increasing levels towards the fully mesenchymal state. Notably, in line with the results of our in vitro models and the pathology analyses, the data from animals demonstrated that the TN amd CD106^+^ E/M states are endowed with the highest metastatic potential (*72*).

### OVOL-mediated epithelialization induces DNA double strand breaks, as well as alters the interactions among the kinases regulating the DNA damage response (DDR)

Stimulation of stress signaling might explain how DTCs survive post cytotoxic treatments (*6, 47*). For example, the stress kinases p38 mitogen activated protein kinase (p38-MAPK) and PERK (*EIF2AK3*) have been implicated in dormancy (*12, 73*). In line with these reports, re-analysis of our RNA sequencing data revealed that OVOL1 overexpression elevated expression of transcripts corresponding to p38delta (*MAPK13*) and *EIF2AK3* (Fig. S6E). We also observed concurrent decreased levels of mRNAs encoding ATM, a master regulator of the cellular response to DNA double-strand breaks (DSBs) (*74, 75*). Because ATM regulates autophagy (*76*) and it also inhibits epithelialization (*77*), we focused on this kinase, along with the two other members of the PIKK family, ATR and DNA-PK, which act as effectors of the DNA damage response (DDR). Notably,

ATM and DNA-PK recognize DSBs whereas ATR responds to single stranded regions. In addition, all three PIKK kinases, as well as p38-MAPK (*78, 79*), phosphorylate the tail of histone variant H2AX, and the phosphorylated form, γH2AX, marks DSBs (*80*).

To stimulate ATM, we treated the inducible derivatives of MDA-MB-231 cells with DOX and later exposed them to a chemotherapeutic agent, carboplatin. Under these conditions we observed phosphorylation of not only ATM and H2AX, but also DNA-PKcs and ATR (Fig. 7A). Notably, overexpression of OVOL1 and OVOL2 reduced ATM’s abundance and, correspondingly, diminished the auto-phosphorylated ATM (pATM). Unexpectedly, overexpression of either OVOL1 or OVOL2 increased phosphorylation of H2AX also in the absence of external stress (meaning, in chemotherapy-naïve cells; Fig. 7A). This, however, did not occur in C1ORF116-overexpressing cells (Fig. S6F). In addition, overexpression of OVOLs increased pAKT, but, as expected, carboplatin partly decreased this effect, in line with the reported ability of DTCs to survive cytotoxic treatments (*7, 81*). Taken together, these observations implied that OVOLs activate a cell survival pathway despite their ability to elevate ROS and induce growth arrest. Along this line, our results also proposed that OVOLs can alter the well-studied kinase interplay that regulates DDR. For example, we noted that OVOL2 suppressed expression of ATM and increased phosphorylation of DNA-PKcs, which might complement the bi-directional inhibitory interactions between DNA-PKcs and ATM (*82, 83*).

**Figure 7:**
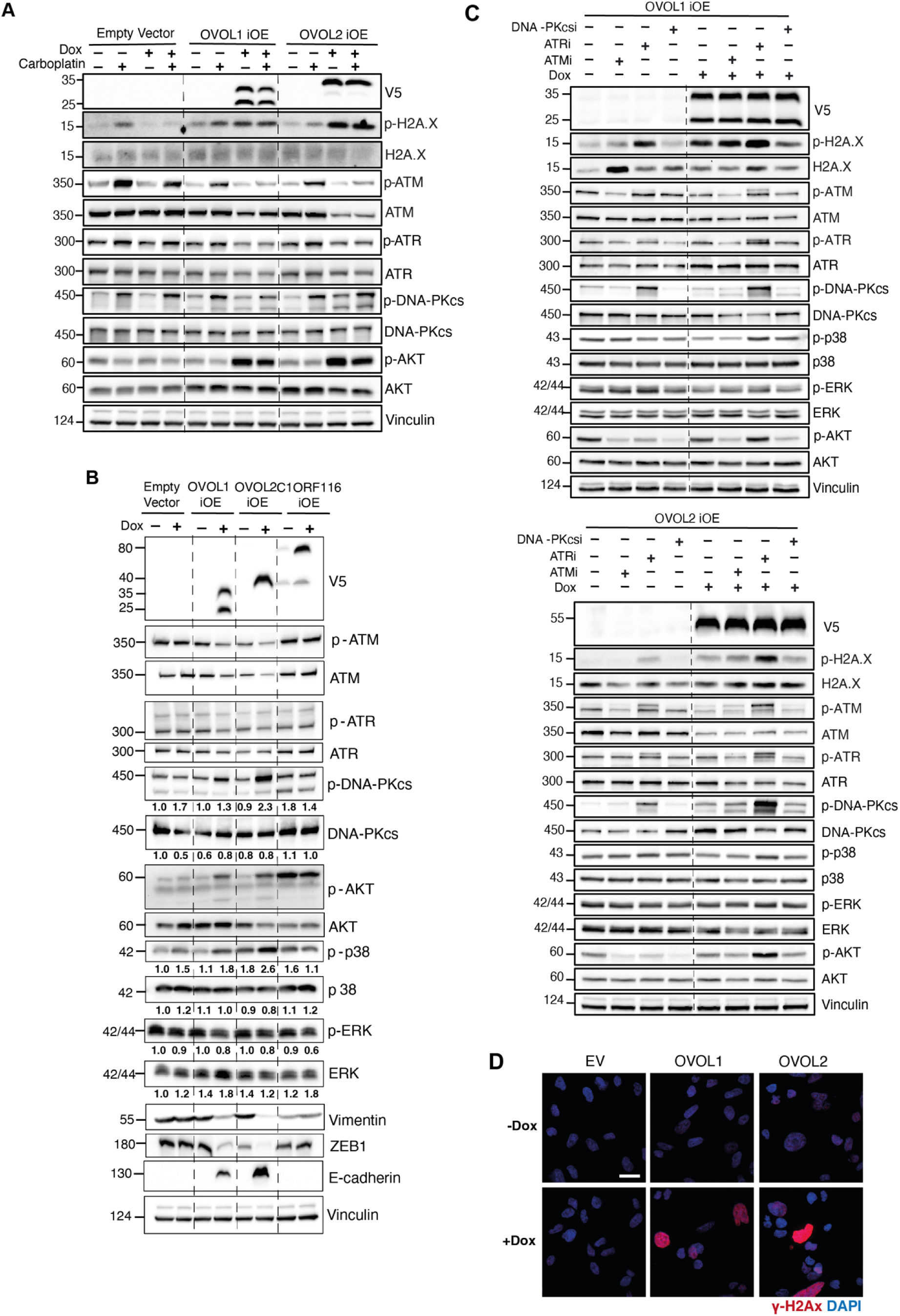
OVOL1 and OVOL2 regulate an interplay among p38-MAPK and the three major kinases involved in the DNA damage response. (**A**) The indicated derivatives of MDA-MB-231 cells, which overexpress V5-tagged OVOL1 and OVOL2, or control (EV) cells, were induced for 3 days with DOX. Thereafter, cells were untreated or treated for 48 hours with carboplatin (15 μM) in the presence of DOX. Next, cell lysates were prepared and subjected to immunoblot analysis that made use of antibodies specific to V5 or to the indicated proteins. Vinculin was used to ensure equal gel loading. (**B**) The indicated derivatives of MDA-MB-231 cells, including cells expressing an inducible allele of C1ORF116 and an empty vector (EV), were untreated or treated for 5 days with DOX. Cell extracts were later cleared and analyzed as in A. (**C**) MDA-MB-231 cells expressing inducible alleles of OVOL1 or OVOL2 were treated for 5 days with DOX and later incubated for 24 hours with the following inhibitors: KU-60019, an ATM inhibitor (ATMi), AZ20, an ATRi, or NU7441, a DNA-PKi. Cell extracts were analyzed as in A. The numbers below specific lanes indicate signal quantification. (**D**) MDA-MB-231 cells overexpressing inducible alleles of OVOL1 and OVOL2 (OVOL1-iOE and OVOL2-iOE, respectively), along with the EV (control) cells, were treated with doxycycline (1 µg/ml) for 72 hours and then they were seeded on glass slides and incubated for 72 additional hours. At the end of the incubation, cells were fixed using paraformaldehyde (PFA; 4%) and immunostained using antibodies specific to the phosphorylated form of histone H2AX (gamma-H2AX). DAPI was used to visualize nuclei. Images were obtained with a Nikon CSU W1-02 spinning disk microscope (x63 magnification). Scale bar, 20 μm.

### Overexpression of OVOL1 and OVOL2 transcriptionally inhibits ATM and catalytically activates DNA-PKcs and p38-MAPK

To better comprehend the impact of OVOL1 and OVOL2 on the DDR, we firstly validated that their gain of function associated with reduced ZEB1 and vimentin, as well as with increased levels of E-cadherin (Fig. 7B). Thereafter, we undertook two approaches: (i) analyzed the effects of OVOL1/2 (in the absence of chemotherapy) on activation of four kinases: the three PIKKs and p38-MAPK (Fig. 7B), which is sensitive to ROS and can phosphorylate H2AX (*74*), and (ii) determined the effects of the respective kinase-specific small molecule inhibitors (Fig. 7C). The first approach validated that overexpression of OVOL1/2 can downregulate ATM and ATR, as well as revealed concomitant activation (phosphorylation) of p38, DNA-PKcs and AKT. Because previous lines of evidence established an inhibitory effect of ATM toward both DNA-PK (*82–84*) and p38 (*85, 86*), the ability of OVOL1/2 to transcriptionally suppress ATM can explain how OVOLs activate DNA-PKcs and p38. Notably, in parallel to the activation of p38, we observed moderate decrease of pERK. These reciprocal trends are consistent with early reports, which inferred that ERK is negatively regulated by p38, hence high p38/pERK activity ratio might herald tumor dormancy (*12, 73*).

Our other approach used the following kinase inhibitors: KU-60019, an ATM inhibitor (ATMi), AZ20, an ATRi and NU7441, a DNA-PKi. In general, the results we obtained (Fig. 7C) further supported the ability of OVOL1/2 to stimulate a dual kinase switch: ATM-to-DNAPK and ERK-to-p38. For example, DNA-PKcs emerged as a major kinase that phosphorylates H2AX in OVOL1/2 overexpressing cells, in line with the observed OVOL1/2-induced decreased expression of ATM and ATR. In addition, using AZ20, an ATRi, we observed activation of DNA-PK and AKT, likely due to relieve of ATR-mediated inhibition of these kinases. The function of ATR, however, remained unclear for the respective inhibitor enhanced rather than reduced pATR. Thus, on the one hand OVOL-induced epithelialization elevates ROS, which activates p38, and on the other hand OVOLs transcriptionally reduce ATM abundance, which activates DNA-PKcs. These biochemical events seem to alter the PIKK interplay and activate a dual kinase switch at DSBs: replacement of ATM by active DNA-PK, along with activating the stress kinase, p38, as a partial substitute for pERK.

### OVOL-mediated epithelialization involves adoption of a **γ**H2AX pattern resembling nucleotide excision repair

To validate an OVOL-induced DNA damage, we analyzed the subnuclear localization of γH2AX. Phosphorylation of this histone variant post exposure to ionizing radiation generates foci serving as DSB markers, but UV irradiation and other stressors induce pan-nuclear staining, which typifies nucleotide excision repair (NER) (*87, 88*). Consistent with the possibility that OVOLs not only enhance γH2AX but also alter its distribution and the identity of the upstream kinases, we noted that control MDA-MB-231 and BT549 cells exhibited faint and rare punctate signals of γH2AX, but DOX-treated cells displayed diffuse pan-nuclear staining (Figs. 7D and S7A). It is noteworthy that pan-nuclear patterns that are caused by localized DNA damage have previously been reported (*87*), but their transient nature and the inferred involvement of ATM and DNA-PK suggest that the underlying mechanism differs from the one activated by OVOL1/2. One potential mechanism could relate to differential degradation of the DDR kinases, as previously reported for ATM and ATR (*89*). Indeed, the application of a proteasome inhibitor and an autophagy blocker raised the possibility that degradation of DNA-PKcs and p38 is controlled by autophagy, whereas ATR and AKT are sorted for proteasomal degradation (Figs. S7B and S7C).

### OVOL-mediated epithelialization involves oxidation of DNA

NER and BER (base excision repair) constantly repair DNA damage caused by ROS and additional mutagens that frequently oxidize the guanine base to produce 8-oxoguanine (which pairs with adenine). Hence, when present in DNA, 8-oxoguanine induces deleterious G > T mutations (*90*). Congruent with the ability of OVOLs to increase ROS levels and alter the DDR, we detected 8-oxoguanine in the cytoplasm of MDA-MB-231 cells posttreatment with DOX (Fig. S7D). Notably, nuclear γH2AX and cytoplasmic 8-oxoguanine displayed a tendency to co- localize in the same cells. In conclusion, the epithelialization of mesenchymal tumor cells involves oxidative stress and adoption of an altered mode of the DDR. This variant involves subnuclear re-distribution of DNA repair sites, re-division of tasks among the involved protein kinases, and possibly altering the way these kinases are controlled by transcription, autophagy and the 26S proteasome.

In summary, we assumed that introducing one of the most characteristic epithelial genes, either OVOL1 or OVOL2, in highly metastatic BC cells will mirror developmental epithelialization and simulate mechanisms that suppress progression of indolent metastases. Here we report that inducible expression of OVOL1/2 in highly metastatic cells preempted their many oncogenes, suppressed their proliferation, as well as inhibited the ability of growth factors to enhance their motility. Further analyses confirmed association of the forced epithelial phenotype with decreased abundance of EMT genes and enhanced abundance of E-cadherin. To unravel the underlying mechanisms, we investigated one of the uncovered OVOL target genes, *C1ORF116*. The respective hormone-controlled adaptor emerged as a putative autophagy receptor that physically regulates two major redox proteins, TRX and GCLC. In line with this, we found that forcing mesenchymal mammary cancer cells to undergo epithelialization associated with lowering glutathione levels and elevating ROS. Yet another unexpected feature emerged from experiments that exposed OVOL-overexpressing cells to carboplatin. Unlike chemotherapy, which activated all three DDR kinases, OVOL overexpression downregulated ATM and ATR, but activated DNA-PKcs, p38-MAPK and AKT. Alongside, OVOL overexpression increased DNA oxidation and DSBs. Taken together, by harnessing C1ORF116-induced putative autophagy and by regulating the redox potential of mesenchymal cells, OVOLs might act as estrogen-inhibitable gatekeepers that prevent exit from breast cancer dormancy while permitting accumulation of deleterious DNA alterations. In aggregate, the observations we report uncovered a hitherto unknown signaling pathway that potentially sustains epithelial phenotypes in development and in pathology (see model in Figure 8).

**Figure 8:**
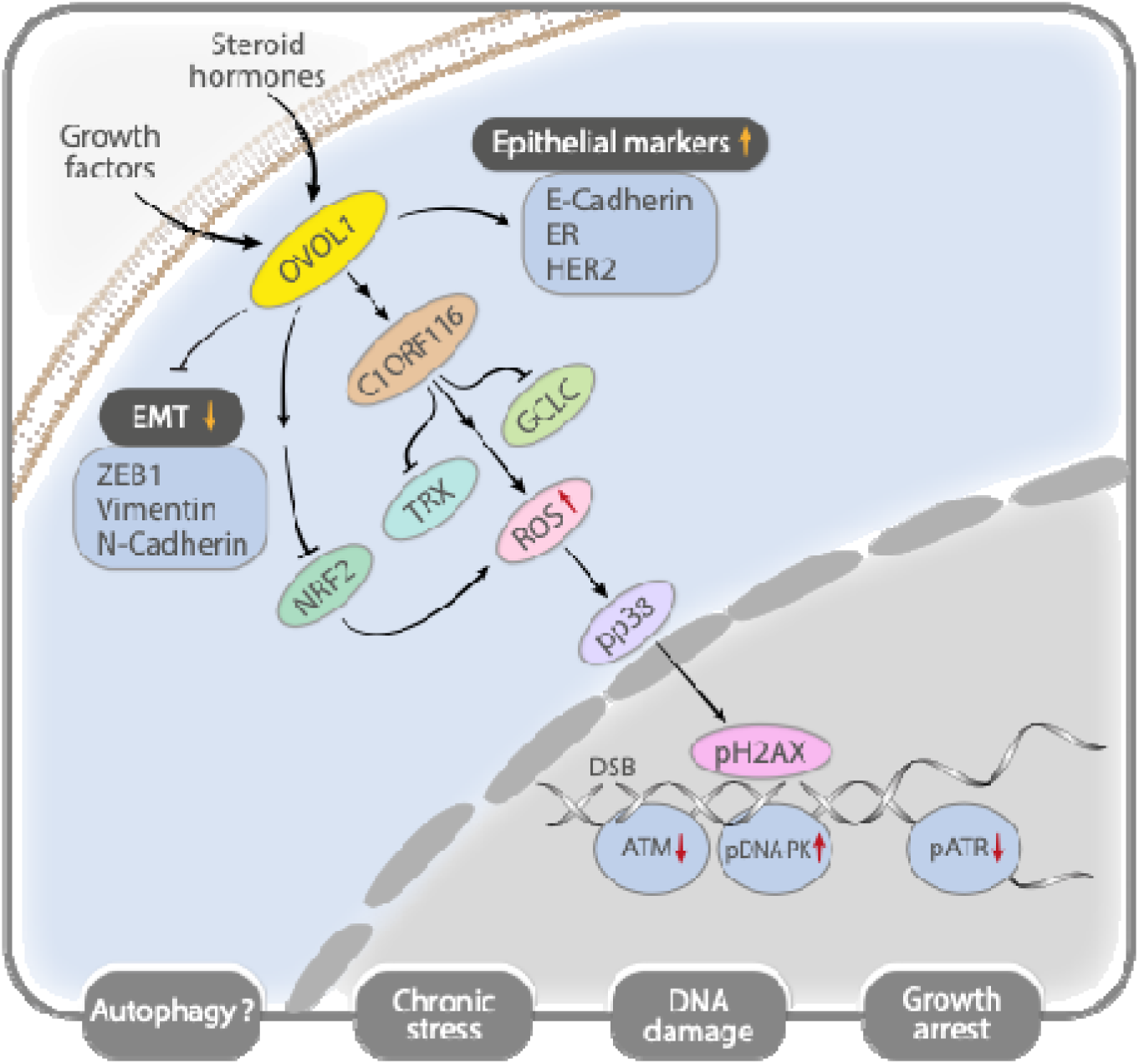
Schematic representation of the inferred OVOL1-regulated signaling pathway that potentially sustains mammary tumor dormancy and epithelial cell identity. Both steroid hormones (such as estrogen) and growth factors (such as EGF) control expression of OVOL1, which represses mesenchymal genes like the ZEB family genes. In parallel, OVOL1 up-regulates epithelial genes that encode for E-cadherin, ER, TROP2 and HER2. Another target is the C1ORF116 gene (chromosome 1, open reading frame 116), which is androgen-regulated and functions as a putative autophagy receptor. C1ORF116 physically binds with GABARAP/LC3 and likely inhibits two major antioxidants, GCLC (glutamate-cysteine ligase catalytic subunit) and thioredoxin (TRX). Thus, C1ORF116 increases the abundance of reactive oxygen species (ROS) and stimulates the p38 Mitogen Activated Protein Kinase (p38-MAPK; also known as Stress Activated Protein Kinase, SAPK). This induces chronic stress and growth arrest, but no accompanied apoptosis, probably because AKT/protein kinase B is also activated. Concurrent with the activation of p38 and AKT in the cytoplasm, transcripts encoding ATM, a nuclear kinase, decrease in OVOL1-expressing epithelial cells. Because ATM normally represses DNA-PKcs, this closely related member of the phosphoinositide 3 kinase related kinase (PIKK) family undergoes stimulation, while the third member of the PIKK family, ATR, undergoes repression. As a result, DNA oxidation is elevated and the tail of histone variant H2AX undergoes phosphorylation at double strand DNA breaks (DSBs). These biochemical events likely herald DNA damage and might explain why awakened tumors commonly display heightened genetic heterogeneity and frequently remain hormone dependent.

## Discussion

An important source of insights into differentiation of the mesenchymal lineage and, by inference, the metastasis cascade, emerged from in-depth understanding of the function of EMT-inducing transcription factors (EMT-TFs), such as ZEB1 (*21, 91*). With the exception of p63 (*92*), we only poorly understand the reciprocal group, MET-TFs, and their involvement in both mammary gland development and BC progression (*93*). Hence, to better understand epithelial lineage differentiation and BC dormancy, we considered several MET-TFs and eventually selected two members of the OVOL group (*17*). During development of the skin and the mammary epithelium, *OVOL1* and *OVOL2* are respectively required for arrest of committed progenitor cells (*94, 95*). Similarly, our results raise the possibility that OVOLs regulate the universal re-epithelialization and growth arrest that DTCs undergo post arrival in distant organs. Mechanistically, OVOL1 directly represses several hub genes, including MYC and ZEB1 (*94, 96*). This explains several attributes of tumor dormancy, such as growth arrest and altered metabolism, along with suppression of EMT. Of note, although we manipulated only single OVOL genes, the engineered cells displayed altered morphological features, along with epithelial markers, such as increased expression of E-cadherin and amphiregulin. In line with these alterations, when using OVOL1/2 overexpressing cells we observed complete inhibition of EGF-induced cell migration, along with partial inhibition of both colony formation (in vitro) and tumor growth (in vivo). Alongside these attributes, we unexpectedly observed increased H_2_O_2_ levels and decreased levels of glutathione, taurine and additional antioxidants. Consistent with these links between redox and tumor dormancy, a recent report attributed the scarcity of muscle colonization by DTCs to sustained oxidative stress and high H_2_O_2_ levels in muscle, as opposed to lung and other organs (*28*). In conclusion, in line with the ability of OVOLs to inhibit antioxidants, the volatile nature of ROS produced by either tumors or their stroma identified a growth arresting metabolic bottleneck that might inhibit exit from dormancy.

Our studies of C1ORF116, a hitherto poorly investigated transcriptional target of OVOL1/2, highlighted yet another feature shared by dormant BC cells: this hormone-inducible adaptor seems to act as an autophagy receptor that supports survival of epithelialized cells under stress conditions. Notably, C1ORF116 has neither recognizable structural domains nor family members. Nevertheless, it undergoes ubiquitination and likely binds with ubiquitin, in similarity to the well-studied autophagy receptor, p62/SQSTM1 (*60*). In line with our results, it has been shown that autophagy and an autophagy-related gene (ATG7) critically support survival of disseminated dormant BC cells (*29*). Likewise, autophagy is utilized as a survival pathway by chemotherapy-treated cells entering a reversible non-proliferative phase that resembles the dormancy state (*97*). In addition, it has been reported that continuous exposure of osteosarcoma cells to insulin can instigate a dormancy state characterized by enhanced autophagy (*98*). According to prevailing models, long-term growth arrest requires elimination of both ROS and misfolded proteins, as well as selective autophagy of mitochondria (*99*). Hence, the herein reported elevation of ROS by the OVOL-C1ORF116 axis was unexpected. Presumably, this might be explained by the dual role played by ROS, namely toxic byproducts of aerobic metabolism, on the one hand, and versatile signaling molecules, on the other hand (*100*).

Apart from identifying OVOL1 and OVOL2 as potential epithelialization promoters and dormancy gatekeepers, which are capable of arresting cell growth and inactivating several antioxidants, our results might provide glimpses into the poorly understood process that allows arrested DTCs to exit from dormancy. Several mechanisms have previously been implicated in this process. They include ageing, remodeling the dormant cell’s niche, post-menopausal obesity and increased neo-vascularization (*6, 47, 101, 102*). Our observations raise yet an additional possibility: by means of downregulating OVOLs, steroid hormones like estrogen and progesterone may abrogate the state of dormancy. Similarly important, our study raises the intriguing possibility that the OVOL-C1ORF-GCLC/TRX signaling pathway functions in conjunction with phosphorylation of histone 2AX and an altered interplay linking p38 and the three major kinases involved in DNA repair (see a model in Figure 8). Conceivably, BC cells in which these pathways are active might slowly accumulate DNA damage and new mutations, till they break a threshold required for escape from dormancy. Stated differently, we propose that unchecked DNA damage that occurs during mass dormancy due to elevated ROS, DNA oxidation and DSBs might increase the genetic heterogeneity of DTCs. This model is consistent with several lines of evidence suggesting that chromosomal instability contributes to the evolution of metastatic disease (*103*). Furthermore, this view is supported by the results of genomic analysis of metastases in autopsies from patients with therapy-resistant BC (*31*). The authors concluded that metastases evolve as communities of clones that keep accumulating new mutations, such that their recurrences are more aggressive and chemo-resistant than the parent tumors. Future studies might refine our model, as well as identify mutational signatures that may underlie escape from the state of dormancy.

## Materials and methods

### Plasmids and antibodies

The cDNA sequences of human OVOL1 (pDONR223 OVOL1-no stop codon, Clone ID: 10848), human OVOL2 (pDONR223 OVOL2-no stop codon, Clone ID: 5098) and human C1ORF116 (pDONR223 C1orf116 no stop codon, Clone ID: 5377) were obtained from CCSB Human ORFeome and cloned into the pLenti6.3/TO/V5-DEST vector to obtain the V5-tagged inducible overexpression vectors. psPAX2 and pMD2.G plasmids were used to produce lentiviral particles from all vectors, along with an Empty Vector control. Promoter reporter clones were obtained from GeneCopoeia. The following antibodies were used: p44/42 MAPK (Erk1/2) (137F5, Cell Signaling Technology #4695), Phospho-p44/42 MAPK (Erk1/2) (Thr202/Tyr204, Cell Signaling Technology #9101), Akt1 (C73H10, Cell Signaling Technology #2938), Phospho-Akt (Ser473, Cell Signaling Technology #4060), Caspase-3 (Cell Signaling Technology #9662), Cleaved Caspase-3 (Asp175, Cell Signaling Technology #9661), Bim (C34C5, Cell Signaling Technology #2933), Phospho-Histone H2A.X (Ser139, Cell Signaling Technology #2577), Histone H2A.X (Cell Signaling Technology #2595), Cyclin B1 (D5C10, Cell Signaling Technology #12231) XP®, Ki-67 (8D5, Cell Signaling Technology #9449), OVOL1 (ProteinTech #14082), OVOL2 (Invitrogen #41620), E-Cadherin (24E10, Cell Signaling Technology #3195), ZEB1 (H-102, Santa Cruz Bio #25388), KLF4 (GKLF/EKLF/LKLF/KLF4/1/2 (F-8, Santa Cruz Bio #166238), Ubiquitin (P4D1, Santa Cruz Bio #8017), P62, SQSTM1 (ProteinTech #18420), DNA-PKcs (3H6, Cell Signaling Technology #12311), Phospho-DNA-PKcs (Ser2056) (E9J4G, Cell Signaling Technology #68716), ATM (D2E2, Cell Signaling Technology #2873), p-ATM (Ser1981) (D6H9, Cell Signaling Technology #5883), ATR (Cell Signaling Technology #2790), p-ATR: (Ser428 Cell Signaling Technology #2853), DNA-PKcs (3H6, Cell Signaling Technology #12311), pDNA-PKcs (Ser2056) (E9J4G, Cell Signaling Technology #68716), alpha Tubulin (Invitrogen #PA5-58711), Glyceraldehyde-3-Phosphate Dehydrogenase (Millipore #MAB374), Estrogen Receptor (Invitrogen, #MA1-310), HER2 (#2165), Beclin-1 (#3738) and Phopsho-Beclin-1 (Ser30; #54101) were from Cell Signaling and p63 (Santa Cruz, #Sc-62686). For immunofluorescence we used Vimentin XP® (D21H3, Cell Signaling Technology #5741), GABARAP+GABARAPL1+GABARAPL2 (EPR4805, Abcam #109364), Thioredoxin (ProteinTech #14999), GCLC (ProteinTech #12601), C1ORF116 (SARG, ProteinTech #14888), NRF2 (Abcam #137550), N-cadherin XP® (D4R1H, Cell Signaling Technology #13116), and V5 (SV5-Pk1, Invitrogen #V9264), E-Cadherin (BD Transduction Laboratories #610404), β- Catenin XP® (D10A8, Cell Signaling Technology #8480), anti-mouse Alexa Fluor 555- conjugated secondary antibody (Thermo Fisher Scientific #A-31570), and anti-rabbit Alexa Fluor 488-conjugated secondary antibody (Thermo Fisher Scientific #A-11008). Fluorescent gelatin (G13186) was from Thermofisher -Aldrich (#V9264).

### Cell lines and cell cultures

MCF-10A, MDA-MB-231 and BT-549 cells were purchased from the American Type Culture Collection. MCF-10A breast cells were cultivated with Dulbecco’s Modified Eagle Medium Nutrient Mixture F-12 (DMEM/F12, Biological Industries, Israel Beit Haemek) supplemented with 5% horse serum (Biological Industries Israel Beit Haemek), 1% Glutamine, 10 ng/ml cholera toxin, 0.5 μg/ml hydrocortisone, 10 μg/ml insulin, 10 μg/ml hydrocortisone and 10 ng/ml human EGF. For the time-course experiments, cells were starved overnight in DMEM/F12 without serum or EGF (starvation medium). MDA-MB-231 and BT549 cells were cultured in RPMI medium (Gibco; Thermo Fisher Scientific, Inc., Waltham MA) supplemented with 10% fetal bovine serum (FCS; Gibco; Thermo Fisher Scientific, Inc., Waltham MA).

### Generation of inducible overexpression cell lines

Inducible overexpression cells were prepared using the pLenti6.3/TO/V5/DEST Gateway destination vector and the pLenti6/TR repressor vector from the ViraPower HiPerform Gateway system (Thermo Fisher Scientific, Inc., Waltham MA). Clones for the cDNA without stop codon for OVOL1 (BC059408), OVOL2 (BC006148), and C1ORF116 (BC000765) were obtained from the CCSB Human ORFeome (hORFeome) collection in Gateway donor plasmids and cloned into the destination vector using LR Clonase II Enzyme Mix (Gateway, Thermo Fisher Scientific). High-efficiency NEB Stable Competent E. coli bacteria (New England Biolabs, Ipswich MA) were transformed with the resulting vectors. Generated pLenti6.3 plasmids and repressor pLenti6/TR plasmid were used to produce lentiviruses using the second-generation packaging systems pMD2.G and psPAX2. Cells were transduced with equal volumes of the overexpression and repressor lentivirus produced in HEK293 cells over 48 hours. Polybrene (10 µg/ml) was used to increase the efficiency of infection. Post-transduction antibiotic selection was performed with blasticidin (5 µg/ml) and Geneticin (1 mg/ml) for MDA-MB-231 cells and Geneticin (0.5 mg/ml) for BT-549 cells.

### Transgenic and xenograft mouse models

Animal studies were approved by the ethics committee of the Weizmann Institute or the Medical University of Vienna and carried out in accordance with guidelines for animal care and protection and protocols approved by the respective authorities. *MMTV^PyMT/+^* mice were described previously (*44*). *Ovol1^tm1a(KOMP)Wtsi^*mice were purchased from MMRRC (Davis, CA, USA) and they were described previously (*43*). Mice were maintained under specific pathogen-free conditions and had access to water and standard rodent diet (V1534, Ssniff, Soest, Germany) *ad libitum*. Beta-galactosidase staining was performed as described previously (*104*). Briefly, the organs of *Ovol1^tm1a(KOMP)Wtsi^* mice, *Ovol1^tm1a(KOMP)Wtsi^* + *MMTV^PyMT/+^* mice, and control littermates were fixed in phosphate buffered saline (PBS) containing 0.02% NP-40, 1% formaldehyde, and 0.2% glutaraldehyde at 4 °C for 2 hours, washed twice with PBS for 20 minutes at room temperature (RT), and incubated overnight at RT under shaking in staining solution containing X-gal (1 mg/ml) in PBS. After staining, samples were washed in PBS as above, postfixed in PBS containing 4% paraformaldehyde and embedded in paraffin blocks. Histological sections were stained with eosin. For tumor growth and metastasis assays, inducible OVOL1 overexpression ZsGreen-labelled MDA-MB-231 cells were incubated with or without doxycycline (1 µg/ml) for 72 hours. For all groups, 1x10^9^ cells were injected into the mammary fat pad of female NOG mice. The indicated mice received DOX (0.5 mg/ml) in the drinking water on the day of cell injection or when a palpable tumor appeared. All groups of animals were monitored for tumor size and well-being. When the tumors approached a size of 600 mm^3^ the experiment was taken to an end and the mice were sacrificed. The tumors were measured and weighed, and the lungs and livers were analyzed for metastatic growths.

### Immunofluorescence staining

8-chamber glass slides were coated with 50µl Cultrex®. MDA-MB-231 cells (5x10^3^/well) were re-suspended in 400µl assay media. The next day, cells were treated with DOX (0.25µg/ml) or left untreated. After 48 hours, cells were treated for 5 min with paraformaldehyde (PFA: 4%) containing sucrose (5%, as fixative) in Triton X-100 (0.2%), and fixed for an additional 25 minutes with the above fixative. Next, cells were washed for 10-min with saline and additional 15-min with saline containing Tween 20 (0.05%). The cells were blocked in IF buffer (130 mM NaCl, 7mM Na_2_HPO4, 3.5 mM NaH_2_PO_4_, 7.7 mM NaN_3_, 0.1% albumin, 0.2% Triton X-100, 0.05% Tween20) containing 5% donkey serum and 5% albumin for 1hr and incubated overnight at 4°C with V5-Tag (D3H8Q) rabbit antibody diluted in the above blocking buffer (1:500; Cell Signaling), for OVOL1/OVOL2 detection, and with Alexa Fluor 488 phalloidin (1:40; Molecular Probes) for F-actin detection. The cells were washed thrice with IF buffer for 15-min, and incubated for 1-hr with donkey ani-rabbit antibody conjugated to Alexa Fluor 647 (1:200; Invitrogen) and mounted with VECTASHIELD mounting medium with DAPI (Vector Laboratories). For F-actin staining only, cells were stimulated with DOX for 6 days before staining was carried out (cells were re-fed after 3 days). Images were captured using a Nikon A1- R confocal laser scanning microscope. To detect NRF2, cover slips were incubated overnight at 4°C with the primary antibody (anti-NRF2; ab137550) diluted in blocking buffer. Cells were washed thrice with saline and subsequently incubated with Alexa Fluor 488-conjugated secondary antibodies, for 60-min in the dark. Cells were washed thrice with saline and incubated for 2-min with Hoechst 33342, washed again and mounted on microscopic slides using mounting media (10 mM phosphate buffer, pH 8.0, 16.6% w/v Mowiol 4–88 and 33% glycerol). A confocal laser-scanning microscope (LSM 800; Carl Zeiss) equipped with an M27 objective lens (Plan Apochromat; Carl Zeiss) was used. Images were captured using a Zeiss Spinning disk confocal microscope and processed using the Zeiss ZEN 3.1 software.

### Cell migration assays

MDA-MB-231 cells were mixed with collagen and then cultured for 48 hours. The left parts of chemotaxis chambers were filled with control media and the right sides with EGF-containing media. Live imaging was performed for 16 hours. The respective rose plots were processed using a dedicated software.

### Cell viability assays

Cell viability was assessed using MTT (3-(4,5-dimethylthiazol-2-yl)-2,5-diphenyltetrazolium bromide). Cells were seeded in 96-well plates. On the next day, they were treated for 72 hours with the indicated drugs. Afterwards, cells were incubated for 3 hours at 37°C with the MTT solution (0.5 mg/ml). The formazan crystals formed by metabolically active cells were dissolved in DMSO and the absorbance was determined at 570 nm.

### Colony formation assays

Cells were seeded in 6-well plates at a density of 1000 cells per well. Media with were refreshed once every 3 days. Following 14 days of incubation, cells were fixed for 20 minutes in ice-cold methanol, followed by staining for 15 minutes at RT with 2% crystal violet. Full-well photos were captured using the EPSON PERFECTION 4870 Photo Scanner (Long Beach, CA, USA). For signal quantification, images corresponding to 5 non-overlapping fields were captured using a light microscope (Olympus Corporation) and quantified using Image J.

### Cell proliferation assays

Cells were seeded in 96-well plates at a density of 1000 cells per well. At the indicated time points, cells were fixed in ice-cold methanol for 20 -min at room temperature, followed by staining for 15 min with 2% crystal violet. Cell growth was quantified by dissolving the cells in a detergent solution and determining light absorbance (590 nm) using a microplate reader. Proliferation assays in 3-D BME made use of Cultrex® growth factor-reduced basement membrane extract (from R&D). MDA-MB-231 cells (2.5x10^3^/well) were cultured in 96-well plates pre-coated with 50µl Cultrex®. Cells were re-suspended in 0.1 ml/well of assay media (RPMI 1640 medium containing Pen-strep, supplemented with 2% FBS and 2% Cultrex®). The next day, cells were treated with Doxycycline (DOX, at a concentration of 0.25 µg/ml) or they were left untreated. Cells were re-fed once every 4 days with assay media with or without DOX. Cell Titer 96 AqueousOne Solution cell proliferation assay kit (Promega) was added to the wells at the indicated time points for 2 hours (*33*). Proliferation was assayed by recording light absorbance at 490nm using an Elisa Plate Reader (Epoch, BioTek).

### Invadopodia formation and gelatin degradation assays

The assay was performed as previously described (*105*). Briefly, 13-mm coverslips were treated with 1 N HCl and coated with poly-l-lysine (50 µg/ml). A gelatin solution (0.2%) was prepared in saline and a 1:10 mixture of Oregon green–labelled gelatin (ThermoFisher)/unlabeled gelatin was warmed to 37°C before addition to the poly-L-lysine–coated plates. Gelatin was crosslinked with 0.01% glutaraldehyde. MDA–MB-231 cells (40,000) were plated on gelatin Alexa Fluor 488–labeled glass coverslips, incubated overnight and fixed in 3.7% paraformaldehyde. Cells were permeabilized with 0.15% Triton X-100, blocked with 1% foetal bovine serum, 1% albumin in saline, and then labelled with rhodamine-phalloidin (Sigma cat# P1951). Images were acquired using an inverted fluorescence microscope (Olympus IX83 60X PlanApo 1,4 NA lens) equipped with an ORCA-Flash 4.0 V2 digital CMOS camera (Hamamatsu Photonics). Invadopodia were identified as actin-rich punctate that colocalized with gelatin degradation areas. Matrix degradation was analyzed by quantifying the mean degraded area in pixels per field using ImageJ and normalized to the number of cells per field.

### Real-time PCR assays, RNA isolation and sequencing

RNA was isolated using the TRIzol reagent (Life Technologies; Thermo Fisher Scientific, Inc., Waltham MA) and cDNA was generated using the qScript cDNA synthesis kit (Quantabio, Beverly, MA). Real-time qPCR relative quantification was performed using the Fast SYBR Green Master Mix (Applied Biosystems). Primer sequences were obtained from the Harvard PrimerBank. Beta-2-microglobulin (B2M) and glyceraldehyde-3-phosphate dehydrogenase (GAPDH) were used as reference genes. Isolated RNA was tested for purity and integrity using NanoDrop quantification and TapeStation analysis, respectively. Pooled libraries were prepared for the selected samples using a modified version of the MARS-Seq protocol for bulk-RNA sequencing. Briefly, the protocol involved the barcoding of samples with oligo dT primers by reverse transcription, pooling the samples for linear amplification and preparation for sequencing. We prepared 4 biological replicates for the cells in culture with 4 consecutive passages. Each sample was sequenced with a depth of at least 5 million reads. Samples were sequenced using the Nextseq 500 platform (75 cycles; Illumina, San Diego CA).

### Evolutionary conservation analysis and LC3 interacting region motif prediction

The analysis of the C1ORF116 sequence conservation was performed using the ConSurf server (https://consurf.tau.ac.il) and 73 homologous sequences from different species. The percentage of residue variety for each amino acid for each of the residues in C1ORF116 was retrieved and projected in a prediction for the 3-D structure of C1ORF116 obtained from Alphafold (https://alphafold.ebi.ac.uk/). For the identification of functional LIRs, we used the LC3 Interacting Region Motifs (https://ilir.warwick.ac.uk/index.php). The complete protein sequence corresponding to C1ORF116 (Uniprot: Q9BW04) and a PSSM score of at least 7 were used.

### Gene expression and pathway analyses

BC gene expression data was obtained from the UCSC Toil RNAseq Recompute Compendium Hub and METABRIC. We retrieved the UCSC Toil data using the R package UCSCXenaTools: R API for UCSC Xena Hubs. This compendium comprises data from The Cancer Genome Atlas (TCGA), the Genotype-Tissue Expression (GTEx), and the Therapeutically Applicable Research To Generate Effective Treatments (TARGET) dataset. Differential expression analysis was performed using the “DESeq2” R package (*106*). The tool EnrichR was used to perform pathway enrichment analysis. Selected genes were used for overrepresentation analysis in the gene expression analysis. Genes with a fold change of at least 1, or -1, and an adjusted p-value smaller than 0.05 were analyzed. Pathway databases used were Elsevier’s Pathway Studio, NCATS BioPlanet, the Molecular Signatures Database and Human Cyc.

### Promoter reporter assays

Promoter containing plasmids were purchased from Genecopoeia (GeneCopoeia, Foster City CA). The assay system used secreted Gaussia Luciferase (GLuc) as the reporter and SEAP (secreted alkaline phosphatase) as the internal control for signal normalization. The reporter system was transfected into HEK293 cells that were treated for 48 hours with the indicated reagents. The medium was collected and processed for measurement of the luminescent signal.

### Immunoblotting analyses

Protein extracts were prepared either from cell lines or from tumors that were excised from mice. Cells were washed in saline and then extracted in RIPA buffer. Proteins were separated using gel electrophoresis and transferred to nitrocellulose membranes. After blocking, membranes were incubated overnight with the indicated primary antibodies, followed by incubation with horseradish peroxidase-conjugated secondary antibodies (1 hour), and treatment with Clarity™ Western ECL Blotting Substrates (Bio-Rad). ECL signals were detected using the ChemiDoc™ Imaging System (Bio-Rad) and images were acquired using the ImageLab Software.

### Coimmunoprecipitation assays

Transfected HEK293T cells were transiently transfected with the C1ORF116-V5 construct using JetPEI (Polyplus-transfection, Illkirch France). Twenty-four hours later, cells were harvested and extracted in RIPA buffer containing protease and phosphatase inhibitors. The extracts (0.5 mg protein) were subjected to coimmunoprecipitation using Pierce Protein A/G Magnetic Beads (Thermo Fisher Scientific) according to the manufacturer’s instructions. Antibodies against V5 and control immunoglobulin G (IgG) were used. Proteins were eluted into SDS gel loading buffer before being subjected to electrophoresis and immunoblotting. For the ubiquitin binding assays, cells were harvested and cell extracts were incubated with ubiquitin-conjugated agarose beads (Enzo Life Sciences, Farmingdale, New York) and naked agarose beads. Electrophoresis and immunoblotting were performed on the pulldown sample.

### ROS production assays

Cells were seeded in 6-well plates (70% confluency) and drugs were added and incubated for 8 hours on the following day. Hydrogen peroxide was determined using 2,7 -dichlorofluorescin diacetate (DCFDA) at a final concentration of 10 mM (diluted in Krebs-Ringer phosphate buffer). Cells were incubated for 30 minutes in the dark at 37°C and 5% CO_2_. After 30 minutes, the cells were washed twice in a fresh Krebs-Ringer phosphate buffer. Finally, the cellular fluorescence signal was recorded using epifluorescence microscopy (Olympus Corporation, Tokyo, Japan) at a wavelength of 500 nm (excitation), and 580 nm emission. Signals were quantified using Image J.

### Mass spectrometry

Cells were rinsed with ice-cold PBS and lysed in RIPA buffer (Thermo Fisher Scientific) supplemented with 1x complete EDTA-free protease inhibitor, 1x PhosSTOP phosphatase inhibitor (both from Roche, Basel, Switzerland), 10 mM NaF, 1 mM Na3VO4, 250 U/ml Benzonase and 10 U/mL RNase-Free DNase (Qiagen). Lysates were incubated on ice for 30 min and cleared by centrifugation. Protein concentrations were determined using the Pierce™ BCA Protein Assay Kit (Thermo Fisher Scientific). For mass-spectromemtry analysis, protein clean-up was performed with 55 µg as input following the automated single-pot solid phase enhanced sample preparation (SP3) workflow adapted from (*107*). Proteins were digested for 16 hours at 37°C with Trypsin in a protease:protein ratio of 1:25. Peptides were vacuum centrifuged to dryness and stored at -20°C. Peptides from 1 µg protein were dissolved in ULC/MS grade water containing 0.1% trifluoracetic acid (TFA) and 2.5% 1,1,1,3,3,3- Hexafluoro-2-propanol (HFIP), followed by sonication for 5 minutes. Peptides were separated using liquid chromatography (Ultimate 3000, Thermo Fisher Scientific) for 100 minutes with a gradient (4-30%) of acetonitrile. The LC system was operated at a flow of 300 nl/min and directly coupled to a MS system (Orbitrap Exploris 480, Thermo Fisher Scientific) via electrospray ionization. MS analysis was performed using data-independent acquisition (DIA). MS1 scans were acquired at a resolution of 120 K covering the range from 350–1400 m/z. Maximum injection time was 45 ms and the automated gain control (AGC) target was set to 3e6. MS2 acquisition was performed using 47 precursor isolation windows of variable width and 1 m/z overlap that covered the range from 400–1000 m/z. Fragment spectra were acquired at a resolution of 30 K and a normalized collision energy of 28% was applied. Maximum injection time was 54 ms and the AGC target was set to 1e6. Raw data were searched against the human proteome with isoforms (downloaded from Uniprot on March 15th, 2022; 79,052 entries) using Spectronaut (v. 17, Diagenode) in directDIA+ mode. The full proteome intensities were median-normalized and Empirical Bayes Statistics for differential enrichment analysis were performed using the limma R package (v. 3.50.3, R version 4.1.3). Missing value imputation was performed using K-Nearest Neighbors (KNN)."

### Statistical analysis

The GraphPad Prism (version 8.0.2) and R programs were used to perform statistical analyses. Sample numbers and other information (mean ± SEM or SD, number of replicates and specific statistical tests) are indicated in the respective figure legends. Differences were considered statistically significant if p<0.05. The Image J, Cell Profiler, R, Image Lab, ReViSP and the IncuCyte S3 software packages were used to perform data analysis.

## Acknowledgments

We thank all members of our laboratory for their insightful comments. In addition, we thank Maxim Itkin, Sergey Malitsky, Noa Wigoda, Shifra Ben-Dor, Hadas Keren-Shaul and Merav Kedmi for performing specific analyses. Special thanks to Prof. Dr. Ursula Klingmüller, Dr. Dominic Helm, and Prof. Dr. Jeroen Krijgsvels for providing mass spectrometry measurement time on a MSCoreSys-funded instrument. This work was performed in the Marvin Tanner Laboratory for Research on Cancer. YY is the incumbent of the Harold and Zelda Goldenberg Professorial Chair in Molecular Cell Biology.

## Funding

This work was supported by the Israel Science Foundation, the European Research Council (ERC), the Israel Cancer Research Fund (ICRF) and the Dr. Miriam and Sheldon G. Adelson Medical Research Foundation.

## Author contributions

DDG and YY conceptualized the experimental design and wrote the manuscript. DDG, SG, RC, ASN, MLUR, AG, MA, AM, NA, TB, ML, AS, NG, LS, NT, H-RM-R, FK, FR, TK, YN, NBN, BW and SD participated in the investigation, performed analytical assays or experiments and analyzing data. DDG, RZ, YEL, NUN and NS performed data curation and formal analysis of genomic and clinical datasets. DDG, RZ and YEL contributed to the visualization of the data. RA and RS provided resources and reviewed the manuscript. GB, SW, SL, OR, CC, ER, YS, MD and YY initiated specific analyses, reviewed results and critically read the manuscript.

## Competing interests

All authors declare that they have no conflicts of interests relevant to the current study.

## Data and materials availability

Transcriptomics data have been submitted to Gene Expression Omnibus (GEO), a public functional genomics data repository (https://www.ncbi.nlm.nih.gov/gds) under the code GSE283235. All materials that are not commercially available will be made available to interested readers. Please contact the lead author, Dr. Yosef Yarden.

## Notes

https://www.ncbi.nlm.nih.gov/geo/query/acc.cgi?acc=GSE283235

### Competing Interest Statement

The authors have declared no competing interest.

## References

1. P. F. Slepicka, A. V. H. Somasundara, C. O. Dos Santos, The molecular basis of mammary gland development and epithelial differentiation. Semin Cell Dev Biol 114, 93–112 (2021).

2. D. P. Cook, B. C. Vanderhyden, Context specificity of the EMT transcriptional response. Nat Commun 11, 2142 (2020).

3. E. Beerling et al., Plasticity between Epithelial and Mesenchymal States Unlinks EMT from Metastasis-Enhancing Stem Cell Capacity. Cell Rep 14, 2281–2288 (2016).

4. L. Bornes et al., Fsp1-Mediated Lineage Tracing Fails to Detect the Majority of Disseminating Cells Undergoing EMT. Cell Rep 29, 2565–2569 e2563 (2019).

5. H. Pan et al., 20-Year Risks of Breast-Cancer Recurrence after Stopping Endocrine Therapy at 5 Years. N Engl J Med 377, 1836–1846 (2017).

6. T. G. Phan, P. I. Croucher, The dormant cancer cell life cycle. Nat Rev Cancer 20, 398–411 (2020).

7. J. Massague, K. Ganesh, Metastasis-Initiating Cells and Ecosystems. Cancer Discov 11, 971–994 (2021).

8. L. Holmgren, M. S. O’Reilly, J. Folkman, Dormancy of micrometastases: balanced proliferation and apoptosis in the presence of angiogenesis suppression. Nat Med 1, 149–153 (1995).

9. C. M. Koebel et al., Adaptive immunity maintains occult cancer in an equilibrium state. Nature 450, 903–907 (2007).

10. S. Malladi et al., Metastatic Latency and Immune Evasion through Autocrine Inhibition of WNT. Cell 165, 45–60 (2016).

11. J. A. Aguirre-Ghiso, Models, mechanisms and clinical evidence for cancer dormancy. Nat Rev Cancer 7, 834–846 (2007).

12. J. A. Aguirre-Ghiso, Y. Estrada, D. Liu, L. Ossowski, ERK(MAPK) activity as a determinant of tumor growth and dormancy; regulation by p38(SAPK). Cancer Res 63, 1684–1695 (2003).

13. D. Barkan et al., Inhibition of metastatic outgrowth from single dormant tumor cells by targeting the cytoskeleton. Cancer Res 68, 6241–6250 (2008).

14. N. Muller-Hermelink et al., TNFR1 signaling and IFN-gamma signaling determine whether T cells induce tumor dormancy or promote multistage carcinogenesis. Cancer Cell 13, 507–518 (2008).

15. J. Hu et al., STING inhibits the reactivation of dormant metastasis in lung adenocarcinoma. Nature 616, 806–813 (2023).

16. A. Dongre, R. A. Weinberg, New insights into the mechanisms of epithelial-mesenchymal transition and implications for cancer. Nat Rev Mol Cell Biol 20, 69–84 (2019).

17. K. Saxena, S. Srikrishnan, T. Celia-Terrassa, M. K. Jolly, OVOL1/2: Drivers of Epithelial Differentiation in Development, Disease, and Reprogramming. Cells Tissues Organs 211, 183–192 (2022).

18. H. Roca et al., Transcription factors OVOL1 and OVOL2 induce the mesenchymal to epithelial transition in human cancer. PLoS One 8, e76773 (2013).

19. J. Chen et al., Comprehensive Analysis of the Expression, Prognosis, and Biological Significance of OVOLs in Breast Cancer. Int J Gen Med 14, 3951–3960 (2021).

20. D. Jia et al., Distinguishing mechanisms underlying EMT tristability. Cancer Converg 1, 2 (2017).

21. I. Pastushenko, C. Blanpain, EMT Transition States during Tumor Progression and Metastasis. Trends Cell Biol 29, 212–226 (2019).

22. D. Jia et al., OVOL guides the epithelial-hybrid-mesenchymal transition. Oncotarget 6, 15436–15448 (2015).

23. X. Dai et al., The ovo gene required for cuticle formation and oogenesis in flies is involved in hair formation and spermatogenesis in mice. Genes Dev 12, 3452–3463 (1998).

24. S. Pasani, S. Sahoo, M. K. Jolly, Hybrid E/M Phenotype(s) and Stemness: A Mechanistic Connection Embedded in Network Topology. J Clin Med 10, (2020).

25. C. Fan et al., OVOL1 inhibits breast cancer cell invasion by enhancing the degradation of TGF-beta type I receptor. Signal Transduct Target Ther 7, 126 (2022).

26. H. Kagawa et al., OVOL1 Influences the Determination and Expansion of iPSC Reprogramming Intermediates. Stem Cell Reports 12, 319–332 (2019).

27. R. Lu et al., Loss of OVOL2 in Triple-Negative Breast Cancer Promotes Fatty Acid Oxidation Fueling Stemness Characteristics. Adv Sci (Weinh), e2308945 (2024).

28. S. B. Crist et al., Unchecked oxidative stress in skeletal muscle prevents outgrowth of disseminated tumour cells. Nat Cell Biol 24, 538–553 (2022).

29. L. Vera-Ramirez, S. K. Vodnala, R. Nini, K. W. Hunter, J. E. Green, Autophagy promotes the survival of dormant breast cancer cells and metastatic tumour recurrence. Nat Commun 9, 1944 (2018).

30. A. La Belle Flynn et al., Autophagy inhibition elicits emergence from metastatic dormancy by inducing and stabilizing Pfkfb3 expression. Nat Commun 10, 3668 (2019).

31. L. De Mattos-Arruda et al., The Genomic and Immune Landscapes of Lethal Metastatic Breast Cancer. Cell Rep 27, 2690–2708 e2610 (2019).

32. D. Barkan et al., Metastatic growth from dormant cells induced by a col-I-enriched fibrotic environment. Cancer Res 70, 5706–5716 (2010).

33. D. Barkan, J. E. Green, An in vitro system to study tumor dormancy and the switch to metastatic growth. J Vis Exp, (2011).

34. K. Weidenfeld et al., Dormant tumor cells expressing LOXL2 acquire a stem-like phenotype mediating their transition to proliferative growth. Oncotarget 7, 71362–71377 (2016).

35. N. Linde, G. Fluegen, J. A. Aguirre-Ghiso, The Relationship Between Dormant Cancer Cells and Their Microenvironment. Adv Cancer Res 132, 45–71 (2016).

36. N. Koons et al., Assessing a Novel 3D Assay System for Drug Screening against OS Metastasis. Pharmaceuticals (Basel) 14, (2021).

37. S. Goossens, N. Vandamme, P. Van Vlierberghe, G. Berx, EMT transcription factors in cancer development re-evaluated: Beyond EMT and MET. Biochim Biophys Acta Rev Cancer 1868, 584–591 (2017).

38. M. Murata et al., OVOL2-Mediated ZEB1 Downregulation May Prevent Promotion of Actinic Keratosis to Cutaneous Squamous Cell Carcinoma. J Clin Med 9, (2020).

39. A. Daemen et al., in Genome Biol. (BioMed Central Ltd, 2013), vol. 14, pp. R110.

40. A. Prat et al., Phenotypic and molecular characterization of the claudin-low intrinsic subtype of breast cancer. Breast Cancer Res 12, R68 (2010).

41. D. Georgopoulou et al., Landscapes of cellular phenotypic diversity in breast cancer xenografts and their impact on drug response. Nat Commun 12, 1998 (2021).

42. C. Curtis et al., The genomic and transcriptomic architecture of 2,000 breast tumours reveals novel subgroups. Nature 486, 346–352 (2012).

43. P. Sun et al., OVOL1 regulates psoriasis-like skin inflammation and epidermal hyperplasia. Journal Investigation Dermatology 141, 1542–1552 (2021).

44. C. T. Guy, R. D. Cardiff, W. J. Muller, Induction of mammary tumors by expression of polyomavirus middle T oncogene: a transgenic mouse model for metastatic disease. Molecular Cell Biology 12, 954–961 (1992).

45. A. Teng, M. Nair, J. Wells, J. A. Segre, X. Dai, Strain-dependent perinatal lethality of Ovol1-deficient mice and identification of Ovol2 as a downstream target of Ovol1 in skin epidermis. Biochim Biophys Acta 1772, 89–95 (2007).

46. J. Joung et al., A transcription factor atlas of directed differentiation. Cell 186, 209–229 e226 (2023).

47. D. Senft, Z. A. Ronai, Adaptive Stress Responses During Tumor Metastasis and Dormancy. Trends Cancer 2, 429–442 (2016).

48. M. S. Sosa, P. Bragado, J. A. Aguirre-Ghiso, Mechanisms of disseminated cancer cell dormancy: an awakening field. Nat Rev Cancer 14, 611–622 (2014).

49. B. Diaz, Invadopodia Detection and Gelatin Degradation Assay. Bio Protoc 3, (2013).

50. M. Blom, K. Reis, P. Aspenstrom, RhoD localization and function is dependent on its GTP/GDP-bound state and unique N-terminal motif. Eur J Cell Biol 97, 393–401 (2018).

51. H. Sun, Q. Li, Z. Song, C. Li, Hypoxia induction of SH2D3A triggers malignant progression of lung cancer. Stem Cell Res 58, 102630 (2022).

52. Q. Liao et al., TROP2 is highly expressed in triple-negative breast cancer CTCs and is a potential marker for epithelial mesenchymal CTCs. Mol Ther Oncol 32, 200762 (2024).

53. C. Rodriguez-Tirado et al., NR2F1 Is a Barrier to Dissemination of Early-Stage Breast Cancer Cells. Cancer Res 82, 2313–2326 (2022).

54. K. Steketee, A. C. Ziel-van der Made, H. A. van der Korput, A. B. Houtsmuller, J. Trapman, A bioinformatics-based functional analysis shows that the specifically androgen-regulated gene SARG contains an active direct repeat androgen response element in the first intron. J Mol Endocrinol 33, 477–491 (2004).

55. Z. He, J. He, K. Xie, KLF4 transcription factor in tumorigenesis. Cell Death Discov 9, 118 (2023).

56. M. Saitoh, Transcriptional regulation of EMT transcription factors in cancer. Semin Cancer Biol 97, 21–29 (2023).

57. D. Rosano et al., Long-term Multimodal Recording Reveals Epigenetic Adaptation Routes in Dormant Breast Cancer Cells. Cancer Discov 14, 866–889 (2024).

58. I. Kalvari et al., iLIR: A web resource for prediction of Atg8-family interacting proteins. Autophagy 10, 913–925 (2014).

59. N. Mizushima, B. Levine, Autophagy in Human Diseases. N Engl J Med 383, 1564–1576 (2020).

60. Z. Liu et al., Ubiquitylation of autophagy receptor Optineurin by HACE1 activates selective autophagy for tumor suppression. Cancer Cell 26, 106–120 (2014).

61. H. F. Aqbi et al., Autophagy-deficient breast cancer shows early tumor recurrence and escape from dormancy. Oncotarget 9, 22113–22122 (2018).

62. Y. Akkoc, N. Peker, A. Akcay, D. Gozuacik, Autophagy and Cancer Dormancy. Front Oncol 11, 627023 (2021).

63. C. Lennicke, H. M. Cocheme, Redox metabolism: ROS as specific molecular regulators of cell signaling and function. Mol Cell 81, 3691–3707 (2021).

64. B. Li et al., Redox Control of the Dormant Cancer Cell Life Cycle. Cells 10, (2021).

65. H. Kumar, R. M. Kumar, D. Bhattacharjee, P. Somanna, V. Jain, Role of Nrf2 Signaling Cascade in Breast Cancer: Strategies and Treatment. Front Pharmacol 13, 720076 (2022).

66. L. H. Saal et al., Recurrent gross mutations of the PTEN tumor suppressor gene in breast cancers with deficient DSB repair. Nat Genet 40, 102–107 (2008).

67. J. Rieckhoff et al., Exploiting Chromosomal Instability of PTEN-Deficient Triple-Negative Breast Cancer Cell Lines for the Sensitization against PARP1 Inhibition in a Replication-Dependent Manner. Cancers (Basel) 12, (2020).

68. S. J. Xu et al., The Specifically Androgen-Regulated Gene (SARG) Promotes Papillary Thyroid Carcinoma (PTC) Lymphatic Metastasis Through Vascular Endothelial Growth Factor C (VEGF-C) and VEGF Receptor 3 (VEGFR-3) Axis. Front Oncol 12, 817660 (2022).

69. H. G. Russnes, O. C. Lingjaerde, A. L. Borresen-Dale, C. Caldas, Breast Cancer Molecular Stratification: From Intrinsic Subtypes to Integrative Clusters. Am J Pathol 187, 2152–2162 (2017).

70. R. N. Batra et al., DNA methylation landscapes of 1538 breast cancers reveal a replication-linked clock, epigenomic instability and cis-regulation. Nat Commun 12, 5406 (2021).

71. Y. Lan, W. Ni, G. Tai, Expression of MUC1 in different tumours and its clinical significance (Review). Mol Clin Oncol 17, 161 (2022).

72. I. Pastushenko et al., Identification of the tumour transition states occurring during EMT. Nature 556, 463–468 (2018).

73. A. C. Ranganathan, L. Zhang, A. P. Adam, J. A. Aguirre-Ghiso, Functional coupling of p38-induced up-regulation of BiP and activation of RNA-dependent protein kinase-like endoplasmic reticulum kinase to drug resistance of dormant carcinoma cells. Cancer Res 66, 1702–1711 (2006).

74. C. S. Schutz, M. B. Stope, S. Bekeschus, H2A.X Phosphorylation in Oxidative Stress and Risk Assessment in Plasma Medicine. Oxid Med Cell Longev 2021, 2060986 (2021).

75. J. H. Lee, T. T. Paull, Cellular functions of the protein kinase ATM and their relevance to human disease. Nat Rev Mol Cell Biol 22, 796–814 (2021).

76. Q. Q. Guo et al., ATM-CHK2-Beclin 1 axis promotes autophagy to maintain ROS homeostasis under oxidative stress. EMBO J 39, e103111 (2020).

77. U. Weyemi et al., The histone variant H2A.X is a regulator of the epithelial-mesenchymal transition. Nat Commun 7, 10711 (2016).

78. Q. B. She, N. Chen, A. M. Bode, R. A. Flavell, Z. Dong, Deficiency of c-Jun-NH(2)-terminal kinase-1 in mice enhances skin tumor development by 12-O-tetradecanoylphorbol-13-acetate. Cancer Res 62, 1343–1348 (2002).

79. A. Kotlyarov et al., Distinct cellular functions of MK2. Mol Cell Biol 22, 4827–4835 (2002).

80. J. Ji et al., Phosphorylated fraction of H2AX as a measurement for DNA damage in cancer cells and potential applications of a novel assay. PLoS One 12, e0171582 (2017).

81. M. P. F. Damen, J. van Rheenen, C. Scheele, Targeting dormant tumor cells to prevent cancer recurrence. FEBS J 288, 6286–6303 (2021).

82. Y. Zhou et al., Regulation of the DNA Damage Response by DNA-PKcs Inhibitory Phosphorylation of ATM. Mol Cell 65, 91–104 (2017).

83. T. Stiff et al., ATM and DNA-PK function redundantly to phosphorylate H2AX after exposure to ionizing radiation. Cancer Res 64, 2390–2396 (2004).

84. B. S. Lee et al., Functional intersection of ATM and DNA-dependent protein kinase catalytic subunit in coding end joining during V(D)J recombination. Mol Cell Biol 33, 3568–3579 (2013).

85. A. Bar-Shira et al., ATM-dependent activation of the gene encoding MAP kinase phosphatase 5 by radiomimetic DNA damage. Oncogene 21, 849–855 (2002).

86. J. Kim, P. K. Wong, Loss of ATM impairs proliferation of neural stem cells through oxidative stress-mediated p38 MAPK signaling. Stem Cells 27, 1987–1998 (2009).

87. B. Meyer et al., Clustered DNA damage induces pan-nuclear H2AX phosphorylation mediated by ATM and DNA-PK. Nucleic Acids Res 41, 6109–6118 (2013).

88. T. M. Marti, E. Hefner, L. Feeney, V. Natale, J. E. Cleaver, H2AX phosphorylation within the G1 phase after UV irradiation depends on nucleotide excision repair and not DNA double-strand breaks. Proc Natl Acad Sci U S A 103, 9891–9896 (2006).

89. A. Cheng et al., ATM loss disrupts the autophagy-lysosomal pathway. Autophagy 17, 1998–2010 (2021).

90. C. Li, Y. Xue, X. Ba, R. Wang, The Role of 8-oxoG Repair Systems in Tumorigenesis and Cancer Therapy. Cells 11, (2022).

91. T. Shibue, R. A. Weinberg, EMT, CSCs, and drug resistance: the mechanistic link and clinical implications. Nat Rev Clin Oncol 14, 611–629 (2017).

92. K. E. Yoh et al., Repression of p63 and induction of EMT by mutant Ras in mammary epithelial cells. Proc Natl Acad Sci U S A 113, E6107–E6116 (2016).

93. J. P. Ng-Blichfeldt, K. Roper, Mesenchymal-to-Epithelial Transitions in Development and Cancer. Methods Mol Biol 2179, 43–62 (2021).

94. M. Nair et al., Ovol1 regulates the growth arrest of embryonic epidermal progenitor cells and represses c-myc transcription. J Cell Biol 173, 253–264 (2006).

95. P. Sun, Y. Han, M. Plikus, X. Dai, Altered Epithelial-mesenchymal Plasticity as a Result of Ovol2 Deletion Minimally Impacts the Self-renewal of Adult Mammary Basal Epithelial Cells. J Mammary Gland Biol Neoplasia 26, 377–386 (2021).

96. G. Tsuji et al., The role of the OVOL1-OVOL2 axis in normal and diseased human skin. J Dermatol Sci 90, 227–231 (2018).

97. E. E. Mowers, M. N. Sharifi, K. F. Macleod, Autophagy in cancer metastasis. Oncogene 36, 1619–1630 (2017).

98. T. Shimizu et al., IGF2 preserves osteosarcoma cell survival by creating an autophagic state of dormancy that protects cells against chemotherapeutic stress. Cancer Res 74, 6531–6541 (2014).

99. A. G. Smith, K. F. Macleod, Autophagy, cancer stem cells and drug resistance. J Pathol 247, 708–718 (2019).

100. S. J. Forrester, D. S. Kikuchi, M. S. Hernandes, Q. Xu, K. K. Griendling, Reactive Oxygen Species in Metabolic and Inflammatory Signaling. Circ Res 122, 877–902 (2018).

101. R. Roy et al., Escape from breast tumor dormancy: The convergence of obesity and menopause. Proc Natl Acad Sci U S A 119, e2204758119 (2022).

102. E. Dalla, A. Sreekumar, J. A. Aguirre-Ghiso, L. A. Chodosh, Dormancy in Breast Cancer. Cold Spring Harb Perspect Med, (2023).

103. S. Turajlic, C. Swanton, Metastasis as an evolutionary process. Science 352, 169–175 (2016).

104. M. Dahlhoff et al., Sebaceous lipids are essential for water repulsion, protection against UVB-induced apoptosis and ocular integrity in mice. Development 143, 1823-1831. Development 143, 1823-1831 (2016).

105. A. Genna et al., Pyk2 and FAK differentially regulate invadopodia formation and function in breast cancer cells. J Cell Biol 217, 375–395 (2018).

106. M. I. Love, W. Huber, S. Anders, Moderated estimation of fold change and dispersion for RNA-seq data with DESeq2. Genome Biol 15, 550 (2014).

107. T. Muller et al., Automated sample preparation with SP3 for low-input clinical proteomics. Mol Syst Biol 16, e9111 (2020).

