## Supplemental Figures for "Inducible re-epithelialization of cancer cells increases autophagy and DNA damage: implications for breast cancer dormancy"

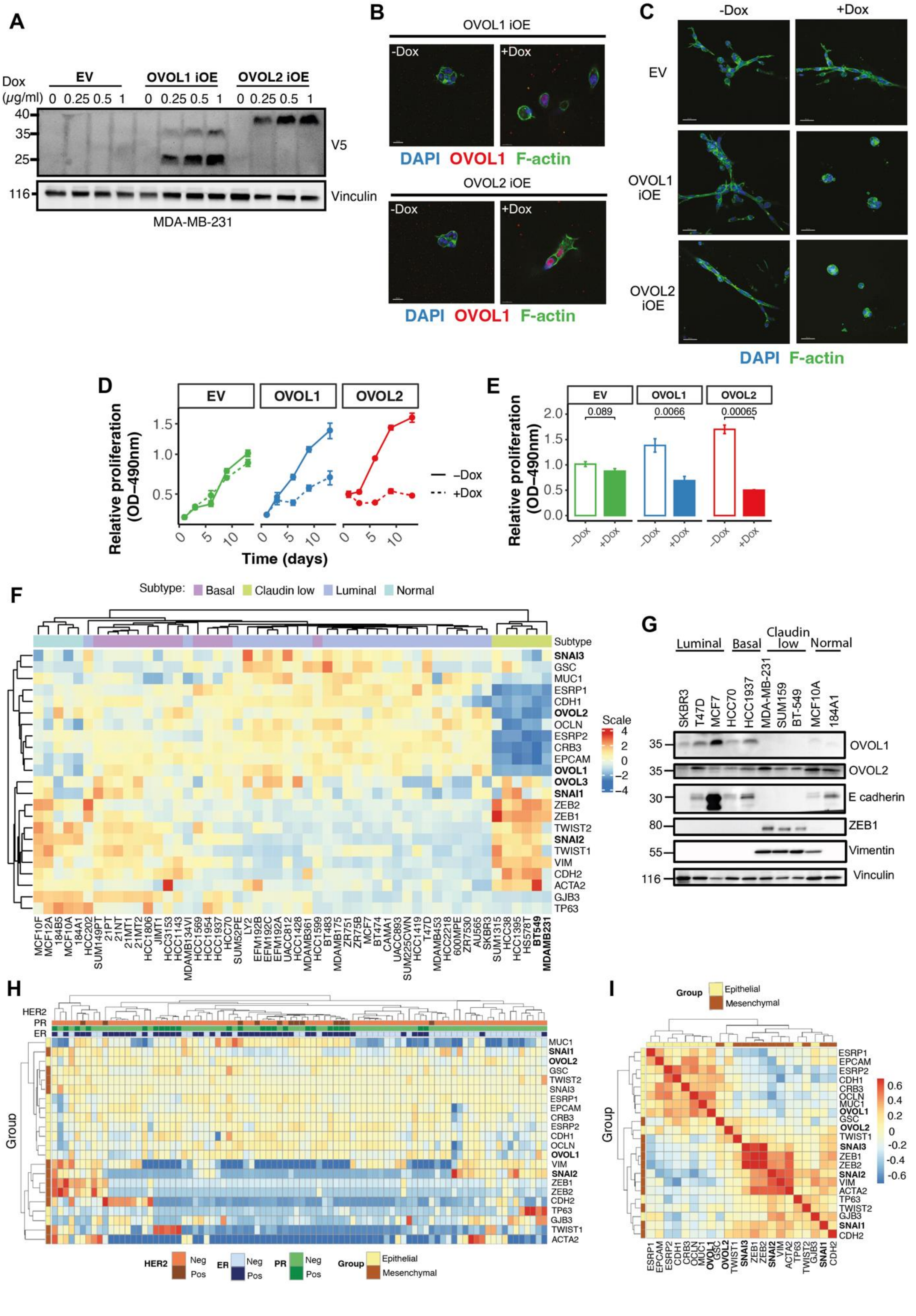

A
Dox (μg/ml)
EV
OVOL1 iOE
OVOL2 iOE
0 0.25 0.5 1
V5
Vinculin
MDA-MB-231
B
OVOL1 iOE
OVOL2 iOE
-Dox
+Dox
DAPI OVOL1 F-actin
C
-Dox
+Dox
EV
OVOL1 iOE
OVOL2 iOE
DAPI F-actin
D
Relative proliferation (OD-490nm)
Time (days)
–Dox
+Dox
E
0.089
0.0066
0.00065
F
Subtype: Basal Claudin low Luminal Normal
Scale
G
Luminal
Basal
Claudin low
Normal
OVOL1
OVOL2
E cadherin
ZEB1
Vimentin
Vinculin
H
HER2
PR
ER
Group
Neg
Pos
Epithelial
Mesenchymal
I
Group
Epithelial
Mesenchymal

**Figure S1: OVOL1 and OVOL2 can induce dormancy in an in vitro model and they are downregulated in mesenchymal breast tumors.** (**A**) MDA-MB-231 inducibly overexpressing (iOE) V5-tagged OVOL1 or OVOL2 were treated for 72 hours with the indicated concentrations of DOX. Cell lysates were prepared and immunoblotted for V5 and vinculin (the loading control). (**B** and **C**) The indicated derivatives of MDA-MB-231 cells were cultured in the 3-D BME matrix and either untreated (-DOX) or treated with DOX (0.25 μg/ml). Shown are immunofluorescence staining photos corresponding to F-actin (green), OVLO1 or OVOL2 (red) after 48 hours of stimulation without or with DOX (B) and F-actin organization after 6 days of stimulation (C; representative confocal images, 40x magnification; scale bar,10 μm). (**D**) Proliferation of untreated or DOX-treated MDA-MB-231 cells (EV, an empty vector control and cells overexpressing inducible OVOL1 and OVOL2). Note that the cells were re-fed once every 4 days. Shown are means and SEM values (bars; n=5). (**E**) Cell proliferation signals from D (day12). The respective p-values are indicated (t-test). Note that the results presented in B through E are representative of 3 biological repeats. (**F**) A dataset of 47 breast cancer cell lines and 5 normal mammary lines was surveyed at the mRNA level for a set of genes relevant to EMT. The uppermost bar indicates the corresponding subtype of BC. Note that OVOL1 and OVOL2 are lowly expressed in claudin-low cell lines characterized by low expression of luminal differentiation markers, along with enrichment for EMT markers. (**G**) Immunoblotting was used to analyze the abundance of OVOL1, OVOL2 and markers of EMT in three claudin-low breast cancer cell lines (MDA-MB-231, SUM159 and BT549) and representatives of the other breast cancer subtypes. (**H**) Gene expression enrichment (log2) of manually curated epithelial and mesenchymal genes in a collection of breast cancer patient-derived tumor xenografts (PDTX). Genes and PDTXs are hierarchically clustered. ER, PR and HER2 status of the PDTXs are shown on the upper colored bars (blue, green and orange). (**I**) Correlation between epithelial and mesenchymal genes across PDTXs (same genes and PDTXs as in H). Genes are hierarchically clustered.

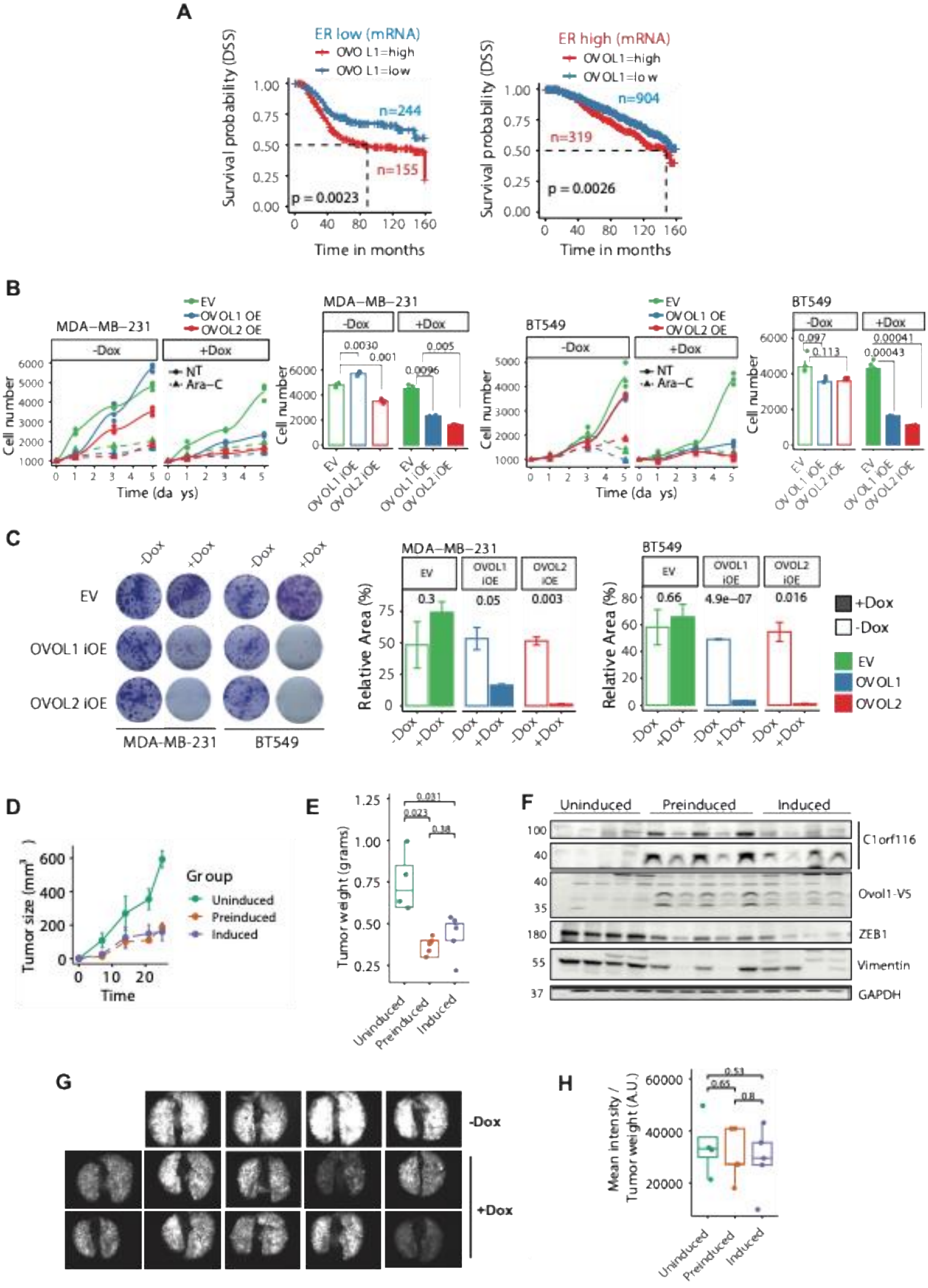
A
ER low (mRNA)
OVOL1=high
OVOL1=low
n=244
n=155
p = 0.0023
ER high (mRNA)
OVOL1=high
OVOL1=low
n=904
n=319
p = 0.0026
Survival probability (DSS)
Time in months
B
MDA−MB−231
BT549
EV
OVOL1 OE
OVOL2 OE
-Dox
+Dox
NT
Ara−C
Cell number
Time (days)
0.0030
0.001
0.005
0.0096
0.097
0.113
0.00041
0.00043
EV
OVOL1 iOE
OVOL2 iOE
C
-Dox
+Dox
EV
OVOL1 iOE
OVOL2 iOE
MDA-MB-231
BT549
Relative Area (%)
0.3
0.05
0.003
0.66
4.9e−07
0.016
OVOL1
OVOL2
D
Tumor size (mm³)
Time
Group
Uninduced
Preinduced
Induced
E
Tumor weight (grams)
0.031
0.023
0.38
F
Uninduced
Preinduced
Induced
C1orf116
Ovol1-V5
ZEB1
Vimentin
GAPDH
G
-Dox
+Dox
H
Mean intensity / Tumor weight (A.U.)
0.53
0.65
0.8

**Figure S2: OVOL1 associates with poor prognosis of patients with breast cancer, but OVOL1-overexpressing cells display reduced tumorigenic growth and no marked effects on metastasis.** (**A**) Disease-specific survival (DSS) probability was calculated using the METABRIC dataset and mRNA data. Prior to analysis, the dataset was stratified according to the status of the estrogen receptor (ER). The number of patients in each group (n) and the respective p-values are indicated. (**B** and **C**) All experiments made use of control (empty vector; EV) MDA-MB-231 and BT549 cells, along with cells inducibly overexpressing (iOE) either OVOL1 or OVOL2. Cells were seeded in 96-well plates at a density of 1,000 cells per well, and on the following day, they were induced (or not) using doxycycline (DOX) to overexpress either OVOL1 or OVOL2. At the indicated time points, cells were fixed in ice-cold methanol for 20 minutes. This was followed by staining for 15 minutes with crystal violet (2%). Cell growth was quantified by dissolving the cells in a detergent solution and determining light absorbance (590 nm). The histograms present results obtained at the last time points. The experiments were repeated thrice, in triplicates (B). Alternatively, cells in 6-well plates were grown for 14 days. The resulting colonies were fixed and stained using crystal violet. Image J was used to quantify the fraction of the area covered by cells. The experiment used 3 biological replicates. Shown are representative images and histograms from one of three experiments (C). (**D**-**H**) The following three groups of female NOG mice (4-5 animals per group) were injected into the mammary fat pad with ZsGreen labeled MDA-MB-231 cells ($1x10^9$) expressing an inducible allele of OVOL1: (i) the mice of the *Preinduced Group* were inoculated with cells pre-treated in vitro for 72 hours with doxycycline (1µg/ml), and DOX was also administered in the drinking water (ad libitum, 200 µg/ml), (ii) the mice of the *Induced Group* received DOX in the drinking water after being inoculated with cells, (iii) the mice of the *Uninduced Group* received no doxycycline, neither before nor after cancer cell inoculation. Tumor size was monitored during the experiment (D), and the primary tumors were weighed at the end of the experiment (E). The collected tumors were used to prepare lysates that were analyzed using immunoblotting with the indicated antibodies (F; each lane corresponds to an animal). In addition, the lungs were retrieved at the endpoint. Shown are images of the fluorescently labeled cells that migrated from the primary tumor (G). The fluorescent signals from the lungs were normalized to the tumor weights at the time of sacrifice (H).

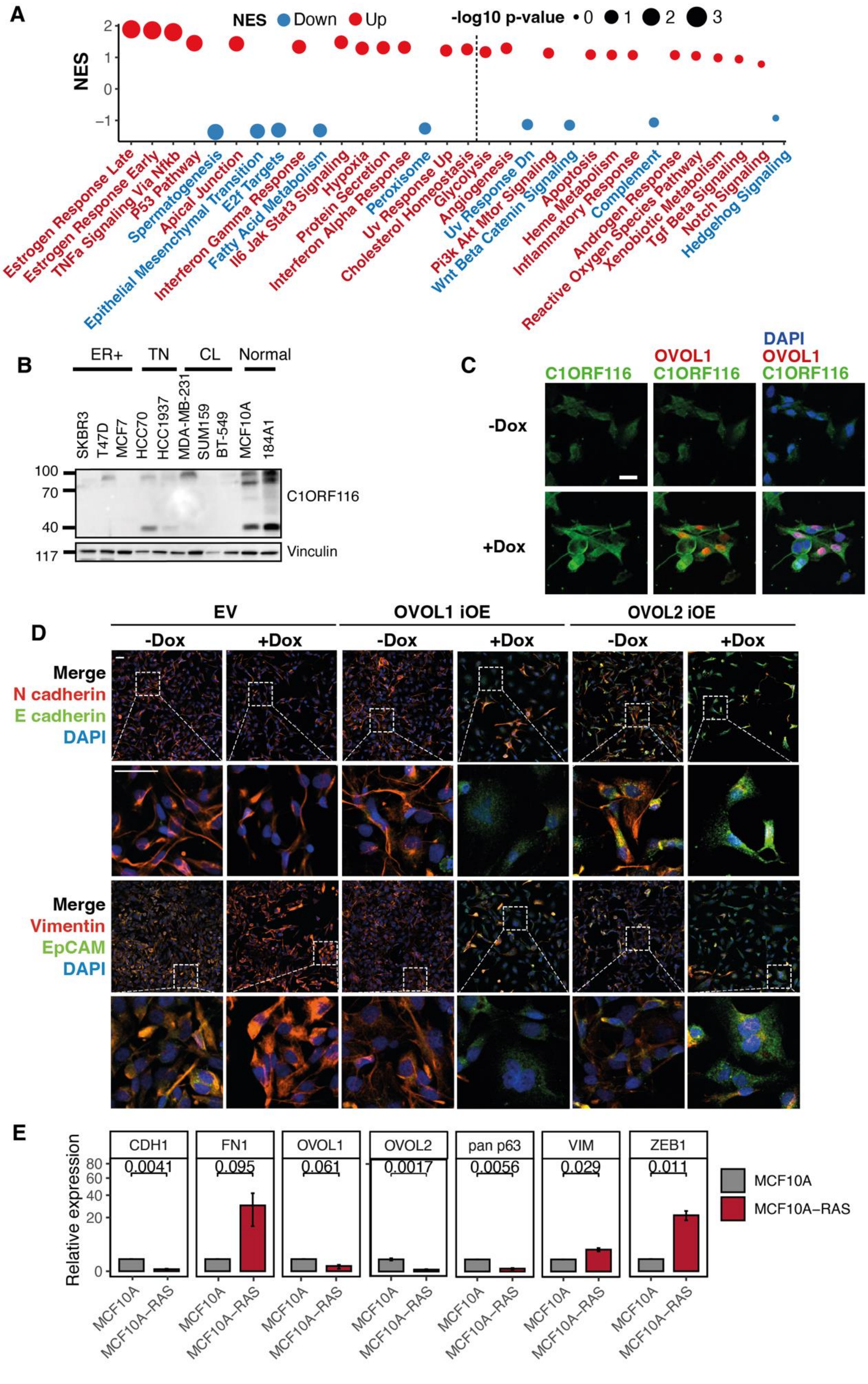
A
NES Down Up
-log10 p-value 0 1 2 3
NES
Estrogen Response Late
Estrogen Response Early
TNFa Signaling Via Nfkb
P53 Pathway
Spermatogenesis
Apical Junction
Epithelial Mesenchymal Transition
E2f Targets
Interferon Gamma Response
Fatty Acid Metabolism
Il6 Jak Stat3 Signaling
Hypoxia
Protein Secretion
Interferon Alpha Response
Peroxisome
Uv Response Up
Cholesterol Homeostasis
Glycolysis
Angiogenesis
Uv Response Dn
Pi3k Akt Mtor Signaling
Wnt Beta Catenin Signaling
Apoptosis
Heme Metabolism
Inflammatory Response
Complement
Androgen Response
Reactive Oxygen Species Pathway
Xenobiotic Metabolism
Tgf Beta Signaling
Notch Signaling
Hedgehog Signaling
B
ER+ TN CL Normal
SKBR3 T47D MCF7 HCC70 HCC1937 MDA-MB-231 SUM159 BT-549 MCF10A 184A1
C1ORF116
Vinculin
C
C1ORF116
OVOL1 C1ORF116
DAPI OVOL1 C1ORF116
-Dox
+Dox
D
EV
OVOL1 iOE
OVOL2 iOE
-Dox +Dox -Dox +Dox -Dox +Dox
Merge
N cadherin
E cadherin
DAPI
Merge
Vimentin
EpCAM
DAPI
E
CDH1 FN1 OVOL1 OVOL2 pan p63 VIM ZEB1
0.0041 0.095 0.061 0.0017 0.0056 0.029 0.011
Relative expression
MCF10A
MCF10A–RAS

**Figure S3: The abundance of epithelial markers is increased following OVOL1 overexpression and their levels decrease following ectopic expression of HRAS.** (**A**) Normalized gene expression data derived from the results shown in Figure 3A were analyzed using Gene Set Enrichment Analysis and the Molecular Signatures Database Hallmarks collection. The dot size is proportional to the statistical significance and the color corresponds to the direction of the enrichment. The hallmarks colored in red are enriched in the OVOL1-OE cells (Up), and the ones in blue are enriched in the EV control cells (Down). Note that the top three up-regulated hallmarks are Estrogen Response Late, Estrogen Response Early and TNF Signaling Via NF-kB. In addition, the top three down-regulated hallmarks are Spermatogenesis, Epithelial Mesenchymal Transition and E2F Targets. (**B**) The indicated human mammary cell lines were grouped into ER-positive breast cancer cells, triple negative (TN), claudin low (CL), and normal cells. Cleared cell extracts were probed for C1ORF116. Vinculin was used to ensure equal gel loading. (**C**) MDA-MB-231 cells expressing an inducible allele of OVOL1 were grown for 72 hours in the absence or presence of doxycycline. Thereafter, the cells were fixed and stained for V5 (OVOL1-V5, red) and C1ORF116 (green). DAPI was used to stain nuclei (blue). Scale bar, 20 µm. (**D**) Three derivatives of BT549 cells (50,000 per well), EV (control), OVOL1-iOE and OVOL2-iOE, were seeded in 8-well Ibidi's slide chambers and then incubated for an additional 72 hours in the presence of DOX. Thereafter, the cells were fixed and subjected to co-immunofluorescence analysis that probed for E-cadherin, N-cadherin, vimentin, and EpCAM. DAPI was used to stain nuclei (blue). Scale bar, 50 µm. (**E**) Messenger RNA was isolated from naïve MCF10A cells or from a stable subline ectopically expressing an oncogenic mutant of human *HRAS*. Quantitative PCR was used to determine the abundance of the indicated transcripts. Shown are results from triplicate assays post normalization to the respective MCF10A levels. The error bars of the MCF10-HRAS levels are 95% confidence levels. All changes are significant within the 95% range, except for fibronectin (FN).

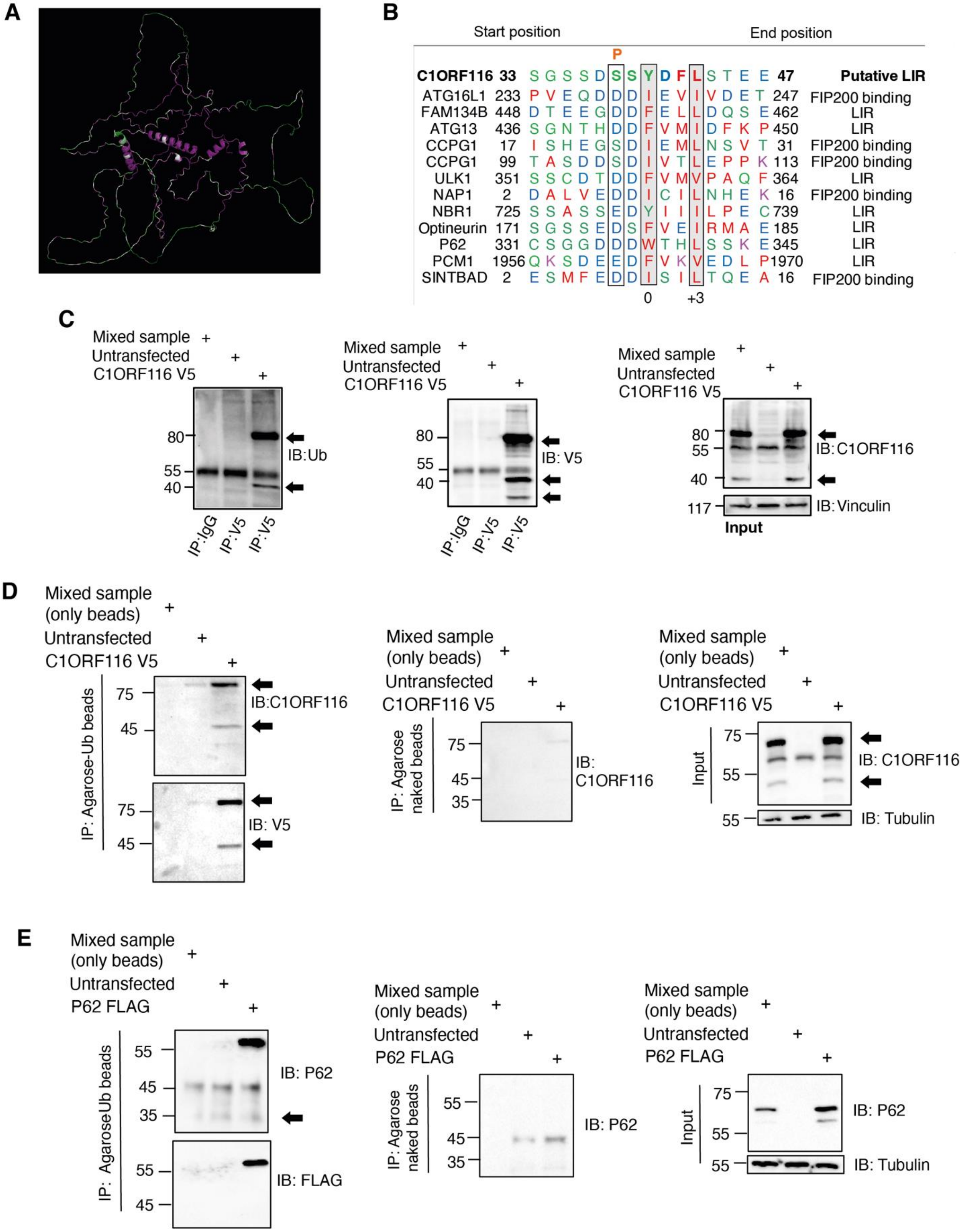


**Figure S4: *C1ORF116* encodes a hormone-regulated protein that undergoes ubiquitination and binds with ubiquitin.** (**A**) The predicted 3D structure of C1ORF116 is shown. Residues with high conservation scores are colored purple, and the residues located within more variable regions are shown in green. Note that the 3D structure was retrieved from Alphafold (https://alphafold.ebi.ac.uk/). (**B**) Sequence alignments comparing the putative LIR of C1ORF116 and the LIRs of several autophagy receptors, including human p62 and optineurin. The gene names and corresponding residue numbers are indicated. The highly conserved acidic residues and the adjacent hydrophobic residues are boxed. The putative phosphorylation site of C1ORF116 is marked ("P"). (**C**) Two 10-cm plates were seeded with HEK293T cells ($1x10^6$). One plate was left un-transfected (control), and the other was transfected with a plasmid encoding a V5-peptide tagged C1ORF116. After 48 hours, cell lysates were prepared and C1ORF116 was pulled-down using protein A/G beads that were pre-incubated with an anti-V5 antibody or with a control immunoglobulin G (IgG). The mixed sample contained lysates from both plates at a 1:1 ratio. The immunoprecipitated (IP) samples were immunoblotted (IB) with an anti-ubiquitin and an anti-V5 antibody. A portion of the lysate (10%) was used as an input control (right panel). An anti-C1ORF116 and an anti-V5 antibody were used to detect the C1ORF116 protein, and vinculin was used to ensure equal gel loading. Note that protein bands corresponding to C1ORF116 are marked (arrowheads) but the immunoglobulin heavy chain (IgH) is unmarked. (**D**) HEK293T cells were treated as in C. One plate was left un-transfected (control), and the other was transfected with a C1ORF116-V5 encoding plasmid. Forty-eight hours later, cell lysates were prepared and ubiquitin-binding proteins were pulled-down using ubiquitin-coated agarose beads. For control, we used naked protein A/G beads. A portion of the lysate (10%) was used as input control. Both input and the pulldown samples were immunoblotted for C1ORF116 or V5. Tubulin was used as a loading control. (**E**) HEK293T cells were treated as in D. One plate was left un-transfected (control), and the other was transfected with a plasmid encoding a FLAG-peptide tagged p62. Forty-eight hours later, cell lysates were prepared and ubiquitin-binding proteins were pulled-down using ubiquitin-coated agarose beads or control protein A/G agarose beads. 10% of the lysate was used as the input control. Both input and the pulldown samples were immunoblotted for p62 and FLAG, as well as for tubulin, the loading control.

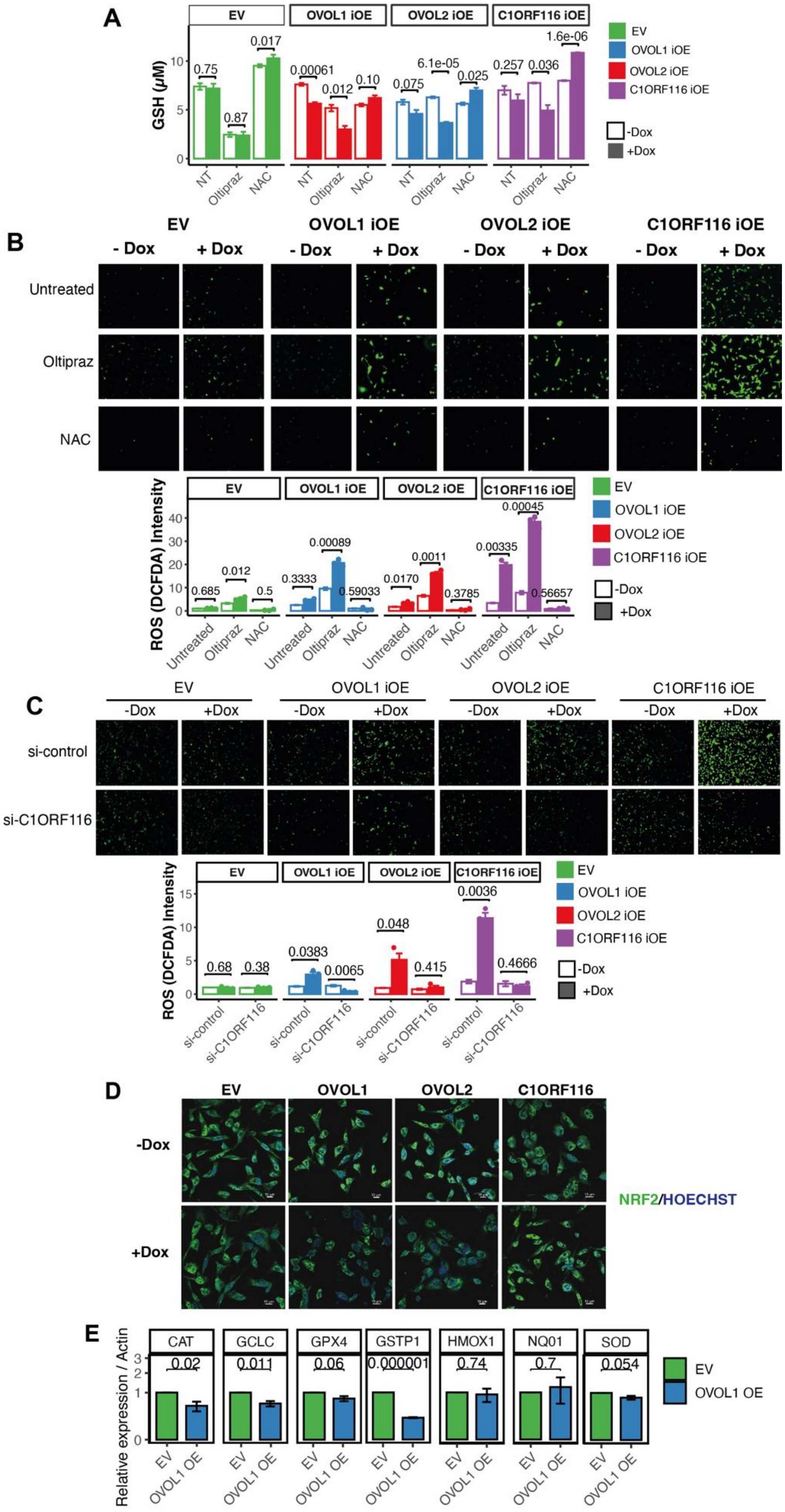
A
EV
OVOL1 iOE
OVOL2 iOE
C1ORF116 iOE
GSH (μM)
NT
Oltipraz
NAC
0.75
0.87
0.017
0.00061
0.012
0.10
0.075
6.1e-05
0.025
0.257
0.036
1.6e-06
-Dox
+Dox
B
- Dox
+ Dox
Untreated
Oltipraz
NAC
ROS (DCFDA) Intensity
0.685
0.012
0.5
0.3333
0.00089
0.59033
0.0170
0.0011
0.3785
0.00335
0.00045
0.56657
C
si-control
si-C1ORF116
0.68
0.38
0.0383
0.0065
0.048
0.415
0.0036
0.4666
D
EV
OVOL1
OVOL2
C1ORF116
NRF2/HOECHST
E
CAT
GCLC
GPX4
GSTP1
HMOX1
NQ01
SOD
0.02
0.011
0.06
0.000001
0.74
0.7
0.054
Relative expression / Actin
EV
OVOL1 OE

**Figure S5: Unlike OVOLs, C1ORF116 promotes proliferation and cannot translocate NRF2 to the cytoplasm of breast cancer cells.** (**A**) The indicated derivatives of BT549 cells were treated with doxycycline (1 µg/ml) for 72 hours. Thereafter, the corresponding cell extracts were subjected to an assay that determined the concentration of reduced glutathione (GSH). Each column represents the mean ± SEM (triplicates) of at least three independent experiments. (**B**) Shown are results of a hydrogen peroxide assay that employed DCFDA (2′,7′-dichlorofluorescin diacetate). Representative images of the inducible BT549 cells overexpressing OVOL1, OVOL2 and C1ORF116 are presented. The images were obtained using epifluorescence microscopy (original magnification X100). Scale bar, 200 µm. The histograms present quantification of hydrogen peroxide levels by densitometric analysis. NAC (N-acetyl-l-cysteine; 10 mM) was used as a ROS scavenger and oltipraz (50 µM) was used to elevate ROS. Values represent mean +SEM (triplicates) of three independent experiments. (**C**) The assay presented in B was repeated except that cells were pre-transfected with siRNA oligonucleotides, either control siRNAs or siRNAs specific to C1ORT116. (**D**) DOX-inducible derivatives of MDA-MB-231 cells were untreated or treated with DOX for 72 hours. Thereafter, the cells were seeded on slides and treated for additional 72 hours. This was followed by fixation and confocal microscopy analysis of NRF2. Hoechst was used to visualize nuclei. Bars, 20 microns. (**E**) RNA was isolated from the indicated DOX-inducible derivatives of MDA-MB-231 cells and after 24 hours it was reverse transcribed to cDNA. Primers specific to the following NRF2 regulated transcripts were employed: NQO1, GCLC, GSTP1, HMOX1, SOD, CAT and GPX4. Shown are the results of three independent experiments. β-actin was used as an internal control. Data represent mean ± SEM of triplicates.

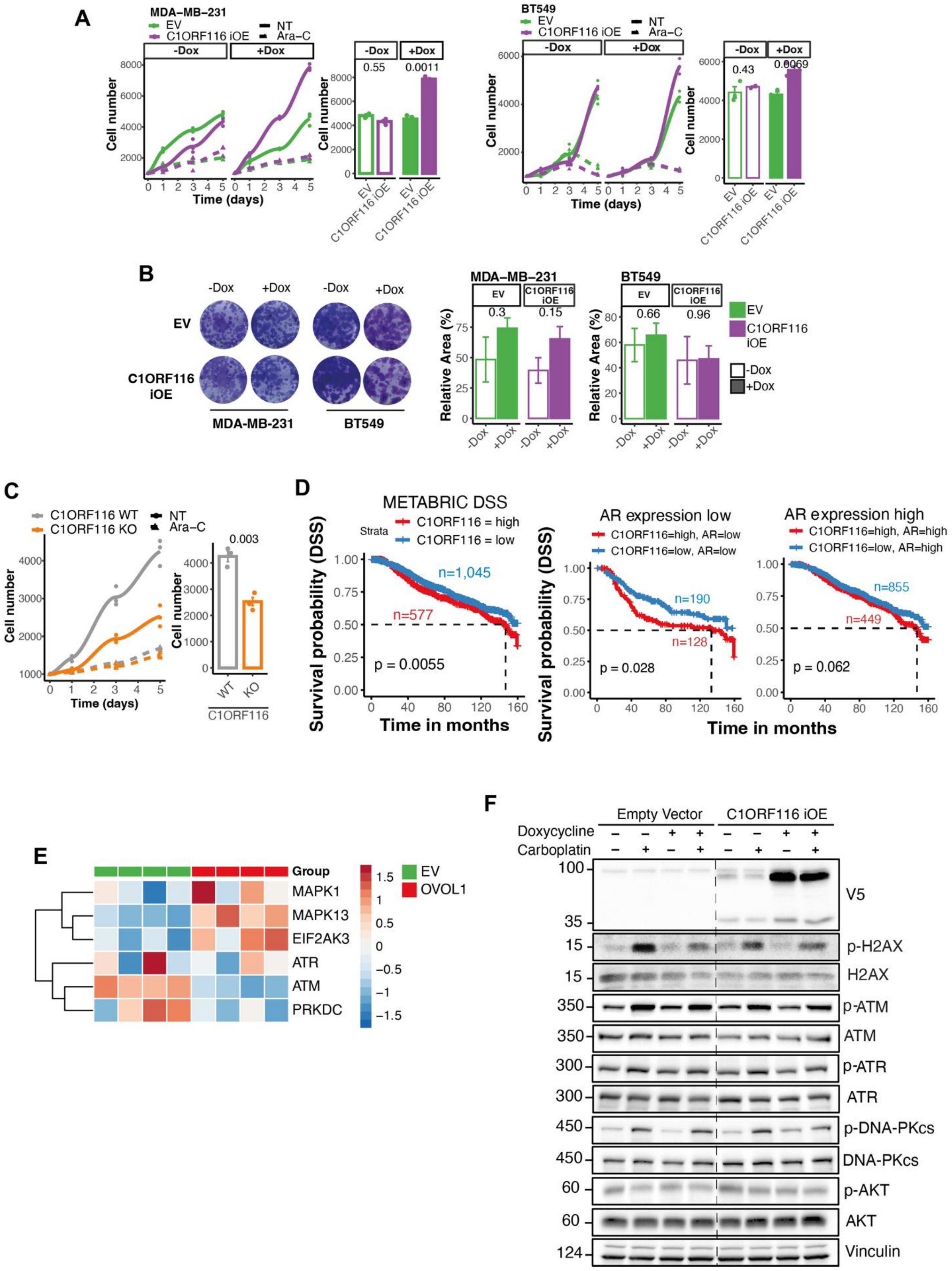
A
MDA–MB–231
EV
C1ORF116 iOE
NT
Ara–C
-Dox
+Dox
Cell number
Time (days)
0.55
0.0011
BT549
0.43
0.0069
B
-Dox
+Dox
EV
C1ORF116 iOE
MDA-MB-231
BT549
Relative Area (%)
0.3
0.15
0.66
0.96
C
C1ORF116 WT
C1ORF116 KO
0.003
WT
KO
C1ORF116
D
METABRIC DSS
Strata
C1ORF116 = high
C1ORF116 = low
n=1,045
n=577
p = 0.0055
Survival probability (DSS)
Time in months
AR expression low
C1ORF116=high, AR=low
C1ORF116=low, AR=low
n=190
n=128
p = 0.028
AR expression high
C1ORF116=high, AR=high
C1ORF116=low, AR=high
n=855
n=449
p = 0.062
E
Group
MAPK1
MAPK13
EIF2AK3
ATR
ATM
PRKDC
EV
OVOL1
F
Empty Vector
C1ORF116 iOE
Doxycycline
Carboplatin
V5
p-H2AX
H2AX
p-ATM
ATM
p-ATR
ATR
p-DNA-PKcs
DNA-PKcs
p-AKT
AKT
Vinculin

**Figure S6: In patients with BC, high C1ORF116 predicts shorter survival of low ER tumors, and in vitro this redox regulator supports cell growth.** (**A**) The following derivatives of MDA-MB-231 (left 2 panels) and BT549 cells (right panels) were used: empty vector (EV) control cells, and cells inducibly overexpressing (iOE) C1ORF116. Cells were pre-treated for 72 hours with doxycycline (1 µg/ml) or vehicle (saline), and then seeded in 96-well plates at a density of 1,000 cells per well. At the indicated time intervals, cells were fixed for 20 minutes in ice-cold methanol and stained for 15 minutes with 2% crystal violet. Cell growth was quantified by dissolving the cells in a detergent solution and determining light absorbance (590 nm). The histogram presents the results obtained at the last time points. We performed a Paired Samples t-test between the C1ORF116-overexpressing cells and the EV control. As reference, we inhibited cell proliferation by treating cells with arabinoside C (Ara-C; 0.3 micromolar). (**B**) Control (EV) MDA-MB-231 and BT549 cells, along with the respective derivatives that inducibly overexpress C1ORF116 (C1ORF116-iOE), were pre-treated for 72 hours with DOX (1 µg/ml). Pre-treated cells (1000 per well) were seeded in 6-well plates and grown for 14 days. The resulting colonies were washed using saline, fixed and stained using crystal violet. Images were captured at high-resolution using a scanner. Image J was used to quantify the fraction of the area covered by cells. The plot shows the fraction of the cell-covered area relative to the whole plate area. The experiment used 3 biological replicates. Shown are representative images and histograms from one experiment. We performed a Paired Samples T-test comparing DOX-treated and vehicle-treated cells. (**C**) Wild type MDA-MB-231 and derivatives lacking C1ORF116 (KO) were seeded in 96-well plates at a density of 1,000 cells per well and grown for the indicated time intervals. Cell proliferation was quantified as in A. The histogram presents the results obtained at the last time points (Student T-test). (**D**) The METABRIC dataset of BC was divided into two groups, high AR (1304 patients) and low AR (318 patients). Next, using the maximally selected rank statistics, each group of patients was stratified into two sub-groups: high expression of C1ORF116 and low C1ORF116. Shown are analyses of disease-specific survival (DSS) in the whole population (left panel) and per group (right panels). Note that patient numbers and p-values are indicated. (**E**) The RNA-sequencing data presented in Figure 3A was analyzed, in quadruplicates, for the expression levels of the indicated transcripts. (**F**) Control MDA-MB-231 (EV), along with the corresponding DOX-inducible C1ORF116-overexpressing cells (C1ORF116-iOE), were treated with doxycycline (1 µg/ml) for 72 hours. Next, we added carboplatin (15 µM) and continued the incubation for additional 48 hours. Cleared cell extracts were subjected to immunoblotting that used the indicated antibodies, including anti-vinculin antibodies, which were used to ensure equal gel loading.

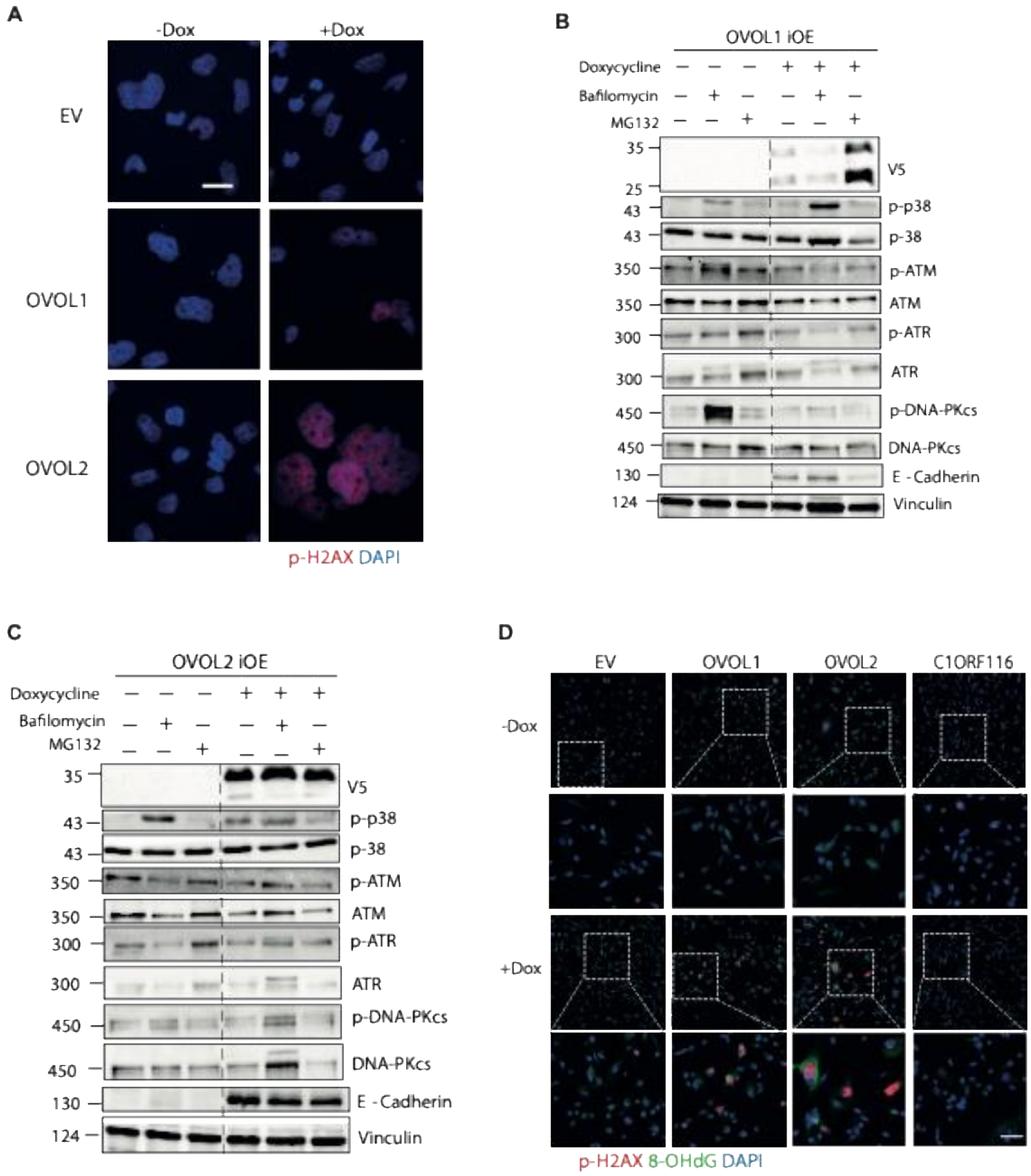


**Figure S7: OVOLs elevate 8-oxoguanine and regulates the PIKK family of DDR kinases.** (A) BT549 cells overexpressing inducible alleles of OVOL1 and OVOL2, along with the EV (control)

cells, were treated as indicated and subjected to immunostaining as in Figure 7D. Scale bar, 20 μm. (**B** and **C**) The indicated DOX-inducible cell lines were treated for 72 hours with DOX (1 μg/ml) or saline. Bafilomycin A1 (20 nM) or MG132 (10 μM) were added 16 or 8 hours (respectively) prior to the end of the 72 hours of incubation. Thereafter, cell extracts were prepared and analyzed as in B using anti-V5 and other antibodies, as indicated. (**D**) DOX-inducible derivatives of MDA-MB-231 cells (50,000; EV, OVOL1-iOE, OVOL2-iOE and C1ORF116-iOE), were seeded in 8-well Ibidi's slide chambers and then incubated for 72 hours in the presence of DOX. Thereafter, the cells were fixed and subjected to staining with antibodies specific to the phosphorylated form of H2AX and to 8-oxoguanine (8-OHdG). DAPI was used to stain nuclei (blue). Scale bar, 50 μm.
